# M4 Muscarinic Receptor Positive Allosteric Modulation Reduces Repetitive Grooming in Male SAPAP3-/- Mice in a Sex-Dependent Manner

**DOI:** 10.64898/2026.11.23.768650

**Authors:** Curran M Johnston, Daniel G Joyner, Yumin Choi, Rebecca D Carden, Michael N May, Margo Wheeler, Pranav Malladi, Laasya Bodducherla, Sreekavya Mekala, Vianne Phan, Casey E. Cryan, Christian O’Reilly, Daniel J. Foster

## Abstract

**Background and Purpose:** Pathological repetitive behaviors in obsessive-compulsive disorder are linked with altered striatal circuitry, and current treatments show variable efficacy and common side effects. We investigated whether the M4 positive allosteric modulator VU0467154 (VU’154) reduces repetitive grooming in SAPAP3-/- mice, and whether M4 receptors expressed in the dorsomedial striatum (DMS) mediate this effect.

**Experimental Approach:** Spontaneous behavior was quantified in male and female SAPAP3-/- mice and wild-type littermates using depth Motion Sequencing (MoSeq) after VU’154 treatment (0.1-3 mg/kg, i.p.). M4 receptors in the DMS were deleted using virally transduced Cre in SAPAP3-/- mice carrying floxed M4 alleles, and receptor deletion was validated by assessing striatal Chrm4 RNA expression.

**Key Results:** SAPAP3-/- mice displayed excessive grooming, hypolocomotion, reduced rearing, and increased inactivity compared to littermate controls. VU’154 reduced grooming in SAPAP3-/- males at a dose that did not significantly affect other active behaviors, while females unexpectedly showed no comparable grooming reduction. Higher doses suppressed active behaviors, including grooming, across genotype and sex. Deletion of M4 from the DMS increased grooming in SAPAP3-/- males but did not prevent VU’154-mediated reductions in grooming.

**Conclusions and Implications:** M4 receptor potentiation selectively reduces excessive grooming in male, but not female, SAPAP3-/- mice, revealing a sex-dependent drug response. M4 receptors in the DMS appear to contribute to baseline grooming regulation in males but were not required for VU’154 efficacy, suggesting that other M4 receptor populations underlie these effects. These findings suggest that sex may be an important consideration in developing M4-targeted treatments for repetitive behaviors.

**Bullet point summary:** *What is already known:*

- SAPAP3-/- mice exhibit robust repetitive grooming associated with disrupted striatal signaling.

*What this study adds:*

- VU0467154 reduced grooming in male, but not female, SAPAP3-/- mice at a dose not affecting other active behaviors
- M4 deletion from the dorsomedial striatum increased grooming in male SAPAP3-/- mice without preventing VU0467154-mediated reductions in grooming

*Clinical significance:*

- These findings support investigating M4 receptors as a target for treatment of repetitive behaviors, with sex-dependent effects warranting careful consideration.

## Introduction

Obsessive-compulsive disorder (OCD) affects approximately 2.3% of US adults at some point in their lifetime (Ruscio, Stein, Chiu, & Kessler, 2010) and is characterized by obsessions and compulsive repetitive behaviors (O’Loghlen, McKenzie, Lang, & Paynter, 2025; Ruscio et al., 2010; Zeidan et al., 2022). These behaviors can significantly disrupt daily functioning, and some repetitive behaviors can cause physical injury (O’Loghlen et al., 2025; Steenfeldt-Kristensen, Jones, & Richards, 2020; Stein et al., 2019). While selective serotonin reuptake inhibitors (SSRIs) are established pharmacological treatments for OCD, their efficacy in treating repetitive behaviors is variable and are often accompanied by side effects (Maneeton, Maneeton, Winichaikul, Kawilapat, Kienngam, & Maneeton, 2024; Skapinakis et al., 2016; Yu et al., 2020; Zhou, Nasir, Farhat, Kook, Artukoglu, & Bloch, 2021). This highlights the need for novel pharmacological approaches to reduce pathological repetitive behaviors without unwanted side effects.

The striatum is the primary input structure of the basal ganglia, and there is considerable evidence of alterations in striatal structure and function in disorders associated with repetitive behaviors (Burton, Longaretti, Zlatanovic, Gomes, & Tonini, 2024; Kim, Lim, & Kaang, 2016; Langen, Bos, Noordermeer, Nederveen, van Engeland, & Durston, 2014; Ting & Feng, 2011). In particular, disruptions in cortico-striatal-thalamo-cortical (CSTC) circuitry have been implicated in the development and expression of these behaviors (Ahmari et al., 2013; Burguiere, Monteiro, Feng, & Graybiel, 2013; Robbins, Banca, & Belin, 2024; Saxena & Rauch, 2000; Ting & Feng, 2011). The SAP90/PSD-95-associated protein 3 (SAPAP3) is a postsynaptic scaffolding protein highly expressed in the striatum (Welch, Wang, & Feng, 2004) and involved in maintaining corticostriatal synaptic function (Welch et al., 2007). Mice lacking SAPAP3 exhibit excessive grooming, accompanied by skin lesions, and the excessive grooming phenotype is rescued by SSRI administration or by restoring SAPAP3 expression in the striatum (Lamothe et al., 2023; Welch et al., 2007). Recent studies have identified alterations in striatal excitability and activity of spiny projection neurons in both direct- and indirect-pathway in SAPAP3 knockout (SAPAP3-/-) mice (Malgady et al., 2023; Piantadosi et al., 2024; Ramirez-Armenta, Alatriste-Leon, Verma-Rodriguez, Llanos- Moreno, Ramirez-Jarquin, & Tecuapetla, 2022), further supporting the involvement of striatal circuitry in repetitive behaviors. These alterations include increased evoked acetylcholine release and attenuated muscarinic modulation of spiny projection neurons in the dorsal striatum (Malgady et al., 2023), providing a rationale for investigating cholinergic modulation in this animal model.

The G protein-coupled muscarinic acetylcholine M4 receptor has emerged as a potential pharmacological target for disorders involving altered striatal function. M4 receptors are highly expressed in the striatum, particularly on direct-pathway spiny projection neurons, where they oppose aspects of dopamine D1 receptor signaling and regulate striatal activity (Jeon et al., 2010; Malgady et al., 2023). M4 receptors are also expressed on cholinergic interneurons, where they function as autoreceptors regulating cholinergic activity, and on presynaptic glutamatergic terminals, where they inhibit corticostriatal transmission (Gonzales, Pare, Wichmann, & Smith, 2013; Pancani, Bolarinwa, Smith, Lindsley, Conn, & Xiang, 2014). Orthosteric agonists bind to highly conserved neurotransmitter binding sites, which makes subtype selectivity difficult to achieve. Positive allosteric modulators (PAMs) bind less conserved sites and can therefore be highly subtype-selective. Several PAMs have been developed that are brain penetrant and highly selective for the M4 subtype of muscarinic acetylcholine receptors (mAChRs) (Bubser et al., 2014; Vuckovic et al., 2023). The M4 PAM VU0467154 (VU’154) has previously been shown to reduce repetitive behaviors in CIN-d and D1CT-7 mouse models of Tourette syndrome (Cadeddu et al., 2024), while another M4 PAM, VU0152100, reduced excessive grooming and other repetitive behaviors in Fmr1 knockout mice, a model of fragile X syndrome (Longo et al., 2023). These findings suggest that M4 PAMs can reduce repetitive behaviors across multiple mouse models.

Although M4 PAMs show promise for suppressing repetitive behaviors, it is unknown whether they do so in a dose-dependent manner without affecting other behaviors. VU’154 has been shown to reduce spontaneous locomotion in mice without impairing other tested aspects of sensorimotor function (Berezovskaia, Lindsley, Fink-Jensen, & Wortwein, 2025). This reduction in spontaneous activity raises the question of whether VU’154 reduces grooming selectively or as part of a broader suppression of motor activity. Conventional measures of grooming and locomotion cannot distinguish selective from nonspecific drug effects. Motion Sequencing (MoSeq) uses machine learning to segment spontaneous behavior into sub-second modules, or syllables, allowing grooming, locomotion, and other observed behaviors to be quantified separately or together (Lin et al., 2024; Wiltschko et al., 2015; Wiltschko et al., 2020). Here, we used MoSeq to characterize spontaneous behavior in male and female SAPAP3-/- mice and assess the dose-dependent effects of VU’154 on excessive grooming and other naturalistic behaviors. The dorsomedial striatum (DMS) appears to play an important role in regulating repetitive behaviors, as optogenetic inhibition of indirect-pathway neurons in the dorsomedial striatum reduces excessive grooming in SAPAP3-/- mice (Ramirez-Armenta et al., 2022). To test if M4 receptors in the DMS regulate repetitive behaviors and mediate VU’154 efficacy, we selectively deleted M4 receptors from this brain region using viral Cre expression in M4 floxed mice. Together, these experiments aim to test the hypothesis that doses of M4 PAMs selectively suppress grooming but not other active behaviors.

## Methods s Materials

### Animals

Experiments utilized both male and female mice that were group-housed and provided with *ad libitum* food and water on a 12-hour light/dark cycle. All experimental procedures were approved by the University of South Carolina’s Institutional Animal Care and Use Committee. SAPAP3-/- (Jax 008733) and control littermates as well as M4fl/fl x SAPAP3-/- mice were used as outlined below (Jeon et al., 2010). Animals were recorded at 3.5 months of age ± 4 weeks. For region-specific M4 deletions, surgery was performed at 2.5 months of age ± 2 weeks, and animals were allowed 1 month for recovery and viral expression prior to behavioral experiments. Animals were transported between the animal facility and the behavioral laboratory in their home cages, covered to reduce stimuli from transport conditions. VU0467154 (VU’154) was generously provided by the Vanderbilt Center for Neuroscience Drug Discovery and was administered intraperitoneally (i.p.) at 10 mL/kg.

### Stereotaxic Surgery

Mice were anesthetized with an isoflurane-oxygen mixture (1.5-2.5% isoflurane) and mounted in a stereotaxic apparatus (Stoelting). Viral vectors were injected using a Hamilton syringe. For viral deletion of M4 receptors, mice received bilateral injections of 0.2-0.3 µL of either Cre recombinase (pENN.AAV5.CamKII 0.4.Cre.SV40, Addgene 105558- AAV5) or control virus (pAAV-CaMKIIa-EGFP, Addgene 50469-AAV5) which was bilaterally injected into the DMS (coordinates from bregma, AP +1.2, ML ±1.3, DV -3.0) at 0.05 µL/min. All mice received post-operative subcutaneous injections of carprofen at 50 mg/kg, with additional doses at 24 and 48 hours post-surgery. At the end of experiments, animals were sacrificed, and brains were removed for RNAscope analysis to confirm viral expression and M4 muscarinic receptor deletion, as outlined below.

### RNAscope

Brains were removed, flash-frozen, sectioned coronally (20 µm) on a cryostat, and assessed using The Advanced Multiplexing RNAscope V2 Assay (Advanced Cell Diagnostics). Briefly, probes (Mm-Chrm1-O1, Mm-Chrm4-C2, Mm-Chat-C3) were conjugated to TSA Vivid 570 (yellow, 1:4000), TSA Vivid 650 (red, 1:5000), and TSA Vivid 520 (green, 1:3000), respectively, and imaged on a Leica Stellaris 5 Confocal Microscope. To quantify M1- and M4-positive cells, the channels were analyzed using QuPath, Fiji, and a custom script (https://doi.org/10.5281/zenodo.23301431; see Supplemental Methods for details).

### Recording Setup

Behavioral recordings were taken in round polyethylene tanks used as open fields, with a 45 cm diameter and 36 cm height. Recordings were obtained using either Azure Kinect depth cameras (Microsoft, Redmond, Washington, United States) or Orbbec Femto Bolt depth cameras (BCH, New York, New York, United States). The latter were operated using the Orbbec wrapper for the Azure Kinect SDK.

### Experimental Timeline

Cohorts were habituated to the open fields for 4 days prior to 30-minute experimental recordings. Baseline recordings were taken the day following habituation. Following baseline recordings, dosage experimental recordings were performed every 48 hours. Each dosage recording day, animals received i.p. injections of either the vehicle (20% Tween 80) or VU0467154 (VU’154), at different dosages (0.1, 0.3, 1, or 3 mg/kg). All animals received each dose, with counterbalanced dosing schedules that were maintained within cohort but randomized between cohorts. Habituation was repeated on days between experimental recordings.

### MoSeq Data Processing

Depth videos were uploaded to Hyperion, the High Computing Cluster of the University of South Carolina, and analyzed using the depth-based Motion Sequencing pipeline (MoSeq2-app, v1.3.1), following published procedures (Lin et al., 2024; Wiltschko et al., 2015). Extraction, quality control, principal component analysis (PCA), κ selection, and final-model selection were performed using the standard functions provided by the pipeline. Extracted depth images were centered and aligned, reduced in dimensionality using PCA, and modeled using an autoregressive hidden Markov model to identify recurring subsecond behavioral syllables.

Four independent MoSeq models were fitted to distinct experimental datasets, consistent with previous MoSeq applications (Wiltschko et al., 2020): one for baseline comparisons between WT and SAPAP3-/- mice, one for their dose-response experiments, one for baseline comparisons between SAPAP3-/- DMS-M4-KO and SAPAP3-/- GFP mice, one for their does- response experiments. Each model was subsequently used to assign frame-by-frame syllable labels to recordings within its corresponding experiment.

Each model was configured to allow a maximum of 100 states. Syllables were ranked by the proportion of frames assigned to each syllable across recordings within each experiment. Only the most frequently expressed syllables accounting cumulatively for 99% of recorded frames were retained for analysis, consistent with a previously published depth-MoSeq application (Song et al., 2025). Syllable usage was calculated as the percentage of recording time assigned to each syllable.

### Syllable Classification

Custom code was used to extract video clips corresponding to individual syllables. Trained scorers viewed these clips and assigned descriptive behavioral labels to each syllable. Classification was repeated using recordings from 6-8 subjects selected semi-randomly to balance experimental groups, sex, and doses where applicable. A behavioral label was assigned when at least 75% of the assessed subject-specific clips for that syllable received the same classification Manual interpretation and grouping of syllables followed approaches used in previous depth-MoSeq studies (Hadjas et al., 2020; Song et al., 2025). Syllables were sorted into behavioral categories: grooming (face/body washing, tail/paw licking, and scratching), active (locomotion, rearing, investigation, and non-locomotor movement), and passive (inactive, freezing, or lacking movement). Behavioral category usage was determined for each by adding together the total usage of all syllables classified in that category for each recording.

### Statistical Analysis

For baseline experiments, two-way ANOVAs were used to determine significance of effects in each behavioral category with genotype and sex as between-subject factors. Where main effects were detected, Fisher’s Least Significant Difference post-hoc test was used to determine significance within genotype and within sex. Pharmacological dose-response experiments used repeated-measures three-way ANOVAs to determine significance in each behavioral category of genotype, sex, and treatment. When significant interactions were found in three-way ANOVAs, analyses were stratified by sex, with separate two-way ANOVAs performed to assess the effects of genotype and treatment. For dose-response experiments, Dunnett’s multiple comparisons were used as a post-hoc test of the two-way ANOVA to determine the significance of changes at each dosage in comparison to vehicle. An α of 0.05 was used to determine significance. All statistical analyses were performed using GraphPad Prism 10.

## Results

### SAPAP3-/- mice exhibit increased grooming and inactivity combined with decreased locomotion and rearing

MoSeq analysis of the baseline recordings of SAPAP3-/- mice (n=18, 9 females, 9 males) and WT control littermates (n = 17, 8 females, 9 males; **Fig. 1A,B**) identified 36 unique high-usage behavioral syllables (most frequently used accounting for cumulative 99% of usage). Syllables (described in **Fig. S1**) were manually classified into 3 groups: grooming behaviors (7 syllables), active non-grooming behaviors (18 syllables), and passive behaviors (11 syllables). Compared with controls, SAPAP3-/- mice showed significant upregulation of 6 syllables, 2 of which were grooming-associated and 4 of which represented passive or inactive behaviors. All 12 syllables downregulated in SAPAP3-/- mice were active, non-grooming behaviors (**Fig. 1C**). Downregulated behaviors were exclusively associated with locomotion and rearing behaviors, which were validated via assessments of tracked velocity and distance traveled **(Fig. S2A,B,E-G)**, and average body height **(Fig. S2C,D)**.

**Figure 1.**
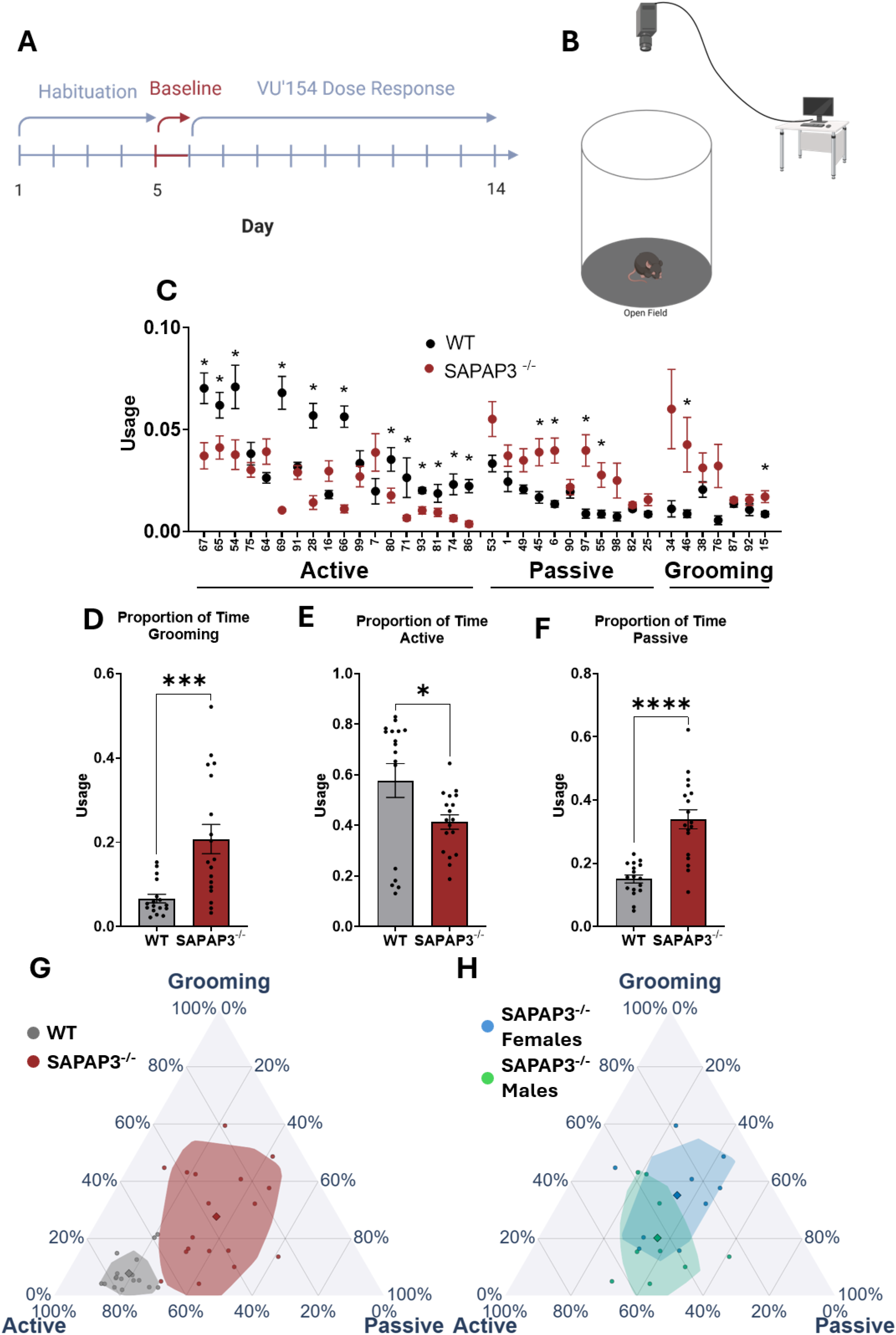
MoSeq analysis of SAPAP3^-/-^ mice as compared to WT control littermates. **A)** 30 min recordings taken after 4 days of habituation with **B)** depth cameras in a circular open field (SAPAP3^-/-^ females/males n=8/9, WT females/males n = 9/9). **C)** Usage comparison of each behavioral syllable between SAPAP3^-/-^ and WT animals identified by MoSeq, organized by broad behavioral category. All highest usage syllables that collectively account for 99% of syllable usage across all recordings. Significance for each syllable determined by independent Mann-Whitney test. **D)** Total usage of all Grooming related syllables compared between WT and SAPAP3^-/-^ mice (Mann-Whitney test p=0.0004). **E)** Total usage of all Active, non-grooming syllables compared between WT and SAPAP3^-/-^ mice (Mann-Whitney test p=0.0380). **F)** Total usage of all Passive syllables compared between WT and SAPAP3^-/-^ mice (Mann-Whitney test p<0.0001). **G-H)** Ternary plot depicting per-recording behavioral usage across the three identified behavioral domains comparing **(G)** SAPAP3^-/-^ (red) and WT (gray) mice and **(H)** SAPAP3^-/-^ male (green) and female (blue) mice. Each point represents a single recording. Stable core calculated as region that appears in 95% of 1000 iterations of randomly selecting 75% of available recordings.

We summed syllable usage within each classified group to identify trends in behavioral differences. Summed syllable usage differed by genotype for grooming (p=0.0005), active (p=0.0236), and passive behaviors (p<0.0001; 3-way ANOVA; **Fig. 1D-G**). Consistent with previous reports (Hadjas et al., 2020), we did not observe any sex differences in any of the baseline behavioral groups (grooming p=0.1046; active p=0.7075; passive p=0.9174; 2-way ANOVA; **Fig. 1H**, **Fig. S1D-F**). MoSeq grooming detection was validated against blinded human scoring in 25 randomly selected videos. For each video, total grooming time was calculated as the proportion of frames scored as grooming by human scorers or assigned to grooming-associated syllables by MoSeq. MoSeq-derived grooming time showed a strong correspondence with manually scored grooming time (R2 = 0.866; **Fig. S3C**.

### VU’154 reduces grooming behavior in male SAPAP3-/- mice

MoSeq analysis was performed for 30 minutes after administration of vehicle or VU’154 (0.1, 0.3, 1, or 3mg/kg; i.p.) to SAPAP3-/- and WT mice (**Fig. 2A,B**). Grooming showed significant main effects of treatment (p=0.0218), genotype (p=0.0188), and sex (p=0.0211), as well as a significant treatment x genotype interaction (p=0.0211; 3-way ANOVA). Active behaviors showed significant main effects of treatment (p<0.0001), and genotype (p<0.0001), but not sex (p=0.1753, 3-way ANOVA). Passive behaviors showed significant main effects of treatment (p<0.0001), genotype (p<0.0001), and sex (p=0.0082; see Supplementary Table 1 for full 3- way and 2-way ANOVA results including F values and degrees of freedom).

**Figure 2.**
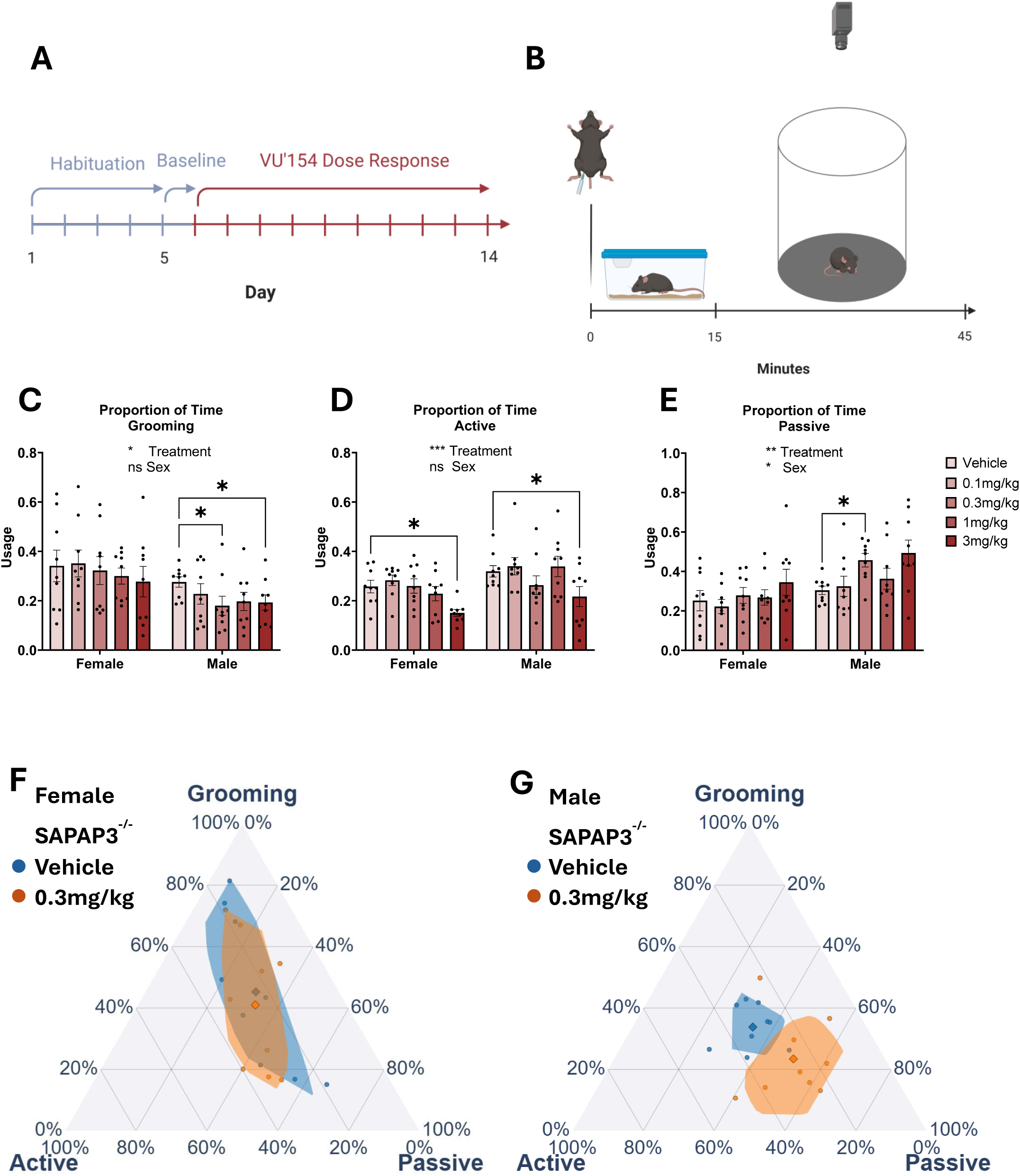
Behavioral analysis of SAPAP3^-/-^ and WT control littermates injected with VU’154. **A)** 30 min recordings taken after 4 days of habituation and baseline recording. Recordings were taken every other day with a counterbalanced dosage schedule used where each animal received each dose once. **B)** Animals were given an i.p. injection and returned to their home cage for 15 minutes prior to recording with depth cameras in a circular open field (SAPAP3^-/-^ females/males n=9/9, WT females/males n = 8/9). **C)** Grooming behavior exhibited main effect of treatment but not sex in SAPAP3^-/-^ animals (2-way ANOVA p=0.0202 and p=0.0893). Reductions in grooming only observed at 0.3mg/kg VU’154 and 3mg/kg VU’154 in male (Dunnett’s post hoc, p=0.0321 and p=0.0464) but not in female (p=0.9611 and p=0.4417). **D)** Active behaviors exhibited main effect of treatment but not sex in SAPAP3^-/-^ animals (2-way ANOVA p=0.0003 and p=0.0728). Reductions were observed at 3mg/kg VU’154 in both male and female animals (Dunnett’s post-hoc, p=0.0383, p=0.0334). **E)** Passive behaviors exhibited main effects in both treatment and sex in SAPAP3^-/-^ (2- way ANOVA p=0.0018 and p=0.0379). Increases in passive behaviors observed at 0.3mg/kg in male but not female SAPAP3^-/-^ mice (Dunnett’s post-hoc, p=0.0162 and p= 0.8915). **F-G)** Ternary plot depicting per-recording behavioral usage across the three identified behavioral domains comparing Vehicle (blue) and 0.3mg/kg dosage trials (orange) within subject in **F)** female SAPAP3^-/-^ mice and **G)** male SAPAP3^-/-^ mice

Due to the significant treatment x genotype interaction, effects on grooming were assessed separately within genotype. In SAPAP3-/- mice, grooming showed a significant main effect of treatment (p=0.0202) but not sex (p=0.0893; 2-way ANOVA). Compared with vehicle, VU’154 reduced grooming in males at 0.3mg/kg (p=0.0321) and 3mg/kg (p=0.0465), but not in females at either dose (p=0.9611 and p=0.4417, respectively; Dunnett’s test; **Fig. 2C**). In WT mice, grooming showed a significant main effect of sex (p=0.0156) and a significant sex x treatment interaction (p=0.0273), but no significant main effect of treatment (p=0.0600; 2-way ANOVA). Compared with vehicle, 3mg/kg VU’154 increased grooming in male mice (p=0.0161) but not in females (p=0.9990, Dunnett’s test; **Fig. S4A**).

In SAPAP3-/- mice, active behaviors showed a significant main effect of treatment (p=0.0003) but not sex (p=0.0728; 2-way ANOVA). Compared with vehicle, 3mg/kg VU’154 reduced active behaviors in both males (p=0.0383) and females (p=0.0334, Dunnett’s test; **Fig. 2D**). In WT mice, active behaviors also showed a significant main effect of treatment (p<0.0001) but not sex (p=0.8969, 2-way ANOVA). Active behaviors were reduced relative to vehicle in males at 1mg/kg (p=0.0009) and 3mg/kg (p=0.0003, Dunnett’s test), but not in females (p=0.8577 and p=0.0944, respectively; Dunnett’s test; **Fig. S4B**).

Passive behaviors in SAPAP3-/- mice showed significant main effects of treatment (p=0.0018) and sex (p=0.0379; 2-way ANOVA). Compared with vehicle, VU’154 increased passive behaviors in males at 0.3mg/kg (p=0.0162) but not in females (p=0.8915; Dunnett’s test; **Fig. 2E**). In WT mice, passive behaviors likewise showed significant main effects of treatment (p=0.0003) and sex (p=0.0358; 2-way ANOVA), with an increase observed at 3mg/kg in males (p=0.0028) but not females (p=0.92016; Dunnett’s test; **Fig. S4C**).

### Selective deletion of M4 receptors from the DMS of SAPAP3-/- mice induces a sex selective increase in the excessive grooming phenotype

We created a model in which M4 is deleted from the dorsomedial striatum (DMS) in SAPAP3-/- mice (DMS-M4-KO) using viral approaches (**Fig. 3A**). In the dorsolateral striatum (DLS) we observed robust expression of M1 and M4 receptor RNA, with M1 RNA expression observed in 93.8% of cells and 48.4% of those M1-positive cells also expressing M4 RNA (**Fig. 3F**). In the DMS, M4 RNA was almost completely ablated in mice receiving AAV encoding Cre-recombinase (0.64% of cells expressing M4 RNA) with little to no change in M1 receptor RNA expression (**Fig. 3E**). M4 RNA expression was lost broadly across the DMS bilaterally but did not extend into the DLS (**Fig. 3B**). Analysis of the number of puncta with M4 or M1 RNA expression in slices with no observed M4 deletion from 7 animals (n=3/4 female/male) showed both subject effects and sex effects in puncta per cell (2-way ANOVA, p<0.0001 and p<0.0001) for expression of both receptors (**Fig. 3E,F**). Total grooming usage in SAPAP3-/- DMS-M4-KO mice showed no effect from M4 deletion but a sex effect (2-way ANOVA, p=0.1389 and p=0.0132) with a trend towards an interaction effect (p=0.0786). An increase in grooming was observed in DMS-M4-KO males compared to GFP males (Fisher’s LSD, p=0.0109) and an increase in grooming in GFP female animals compared to GFP males (p=0.0024) with comparable grooming observed in both female groups and male DMS-M4-KO animals (**Fig. 4C**). Active behaviors showed a sex but not an M4 deletion effect (2-way ANOVA, p=0.0077 and p=0.0621) with a decrease in active behaviors in DMS-M4-KO males compared to GFP males (Fisher’s LSD, p=0.0112) and in GFP female animals compared to GFP males (p=0.0024; **Fig. 4D**). Passive behaviors exhibited neither an M4 deletion or sex effect (2-way ANOVA, p=0.7398 and p=0.6442; **Fig. 4E**).

**Figure 3.**
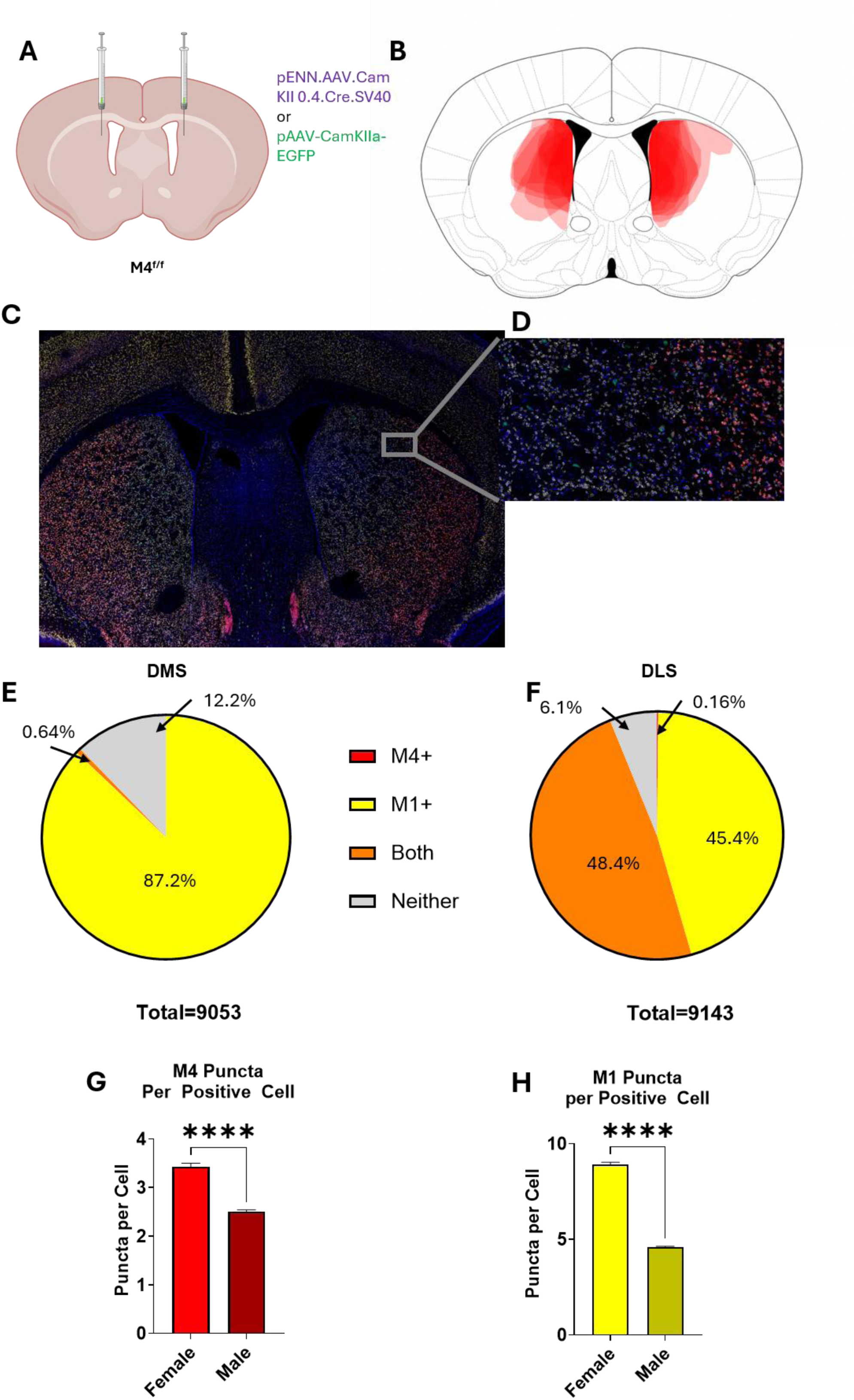
Deletion of M4R from DMS in SAPAP3^-/-^ animals. **A)** Diagram depicting viral injections. Animals were injected bilaterally with 0.2-0.3uL of either pENN.AAV.CamKII 0.4.Cre.SV40 or pAAV-CamKIIa-EGFP. **B)** Diagram depicting overlays of M4 deletion area, with fluorescent images fitted to brain atlas. **C)** Representative RNAscope image of M4 deletion (blue = DAPI, yellow = M1, red = M4). **D)** Closeup showing M4 deletion as a clean delineation between pENN.AAV.CamKII 0.4.Cre.SV40 expression area. **E-F)** Percent of cells expressing M4 and/or M1 mRNA in both the DMS **(E)** and **(F)** DLS. **G-H)** M4 **(G)** and M1**(H)** RNA puncta expression counts per positive cell in the dorsal striatum in both female (n=3) and male (n=4) animals exhibit main effects of both sex and animal (2-way ANOVA, p<0.0001 and p<0.0001).

**Figure 4.**
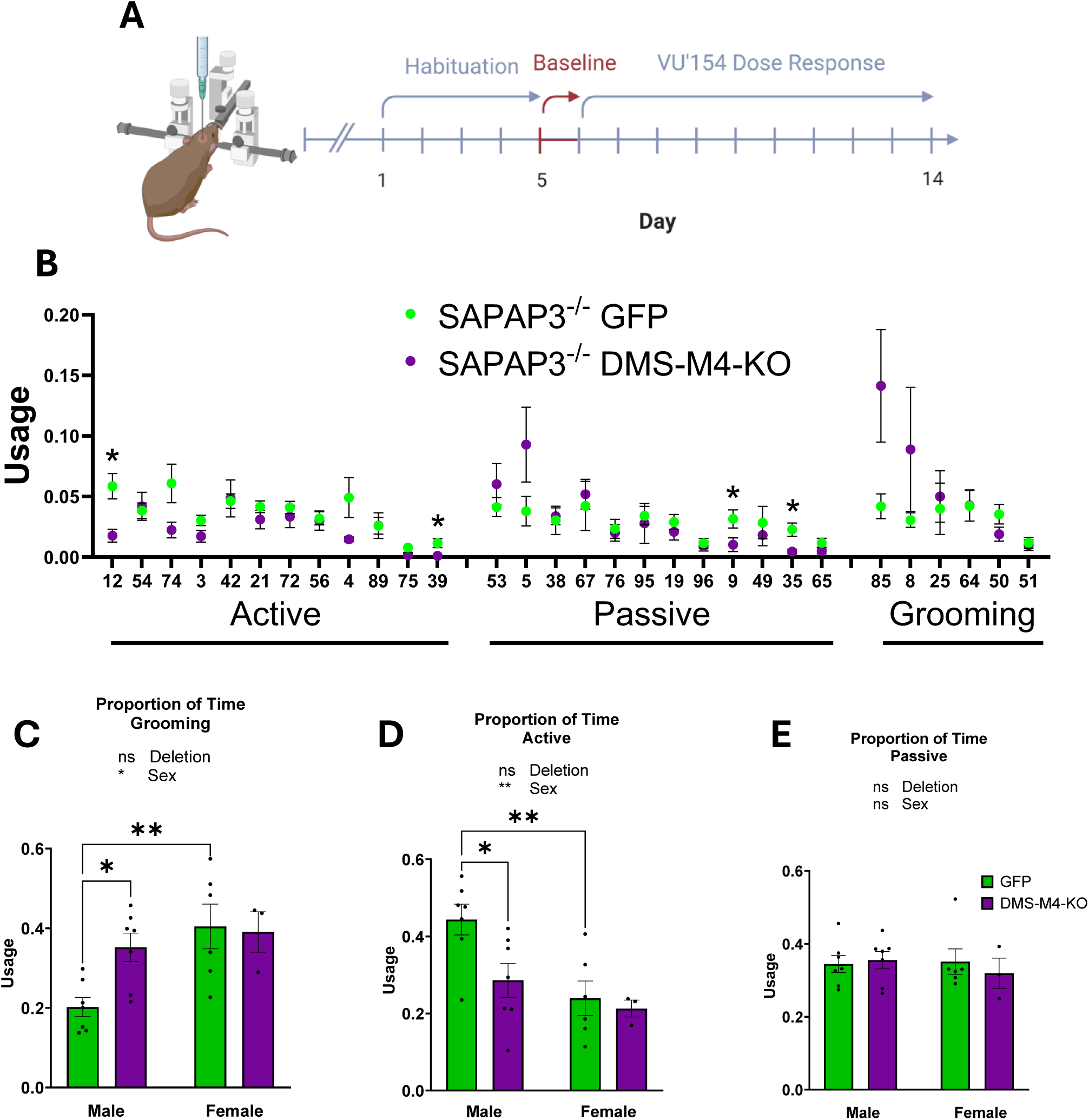
Behavioral analysis of SAPAP3^-/-^ DMS-M4-KO. **A)** Animals received viral injections 1 month prior to experimental timeline. 30-minute recordings taken after 4 days of Habituation (n=7/3 DMS-M4-KO males/females and n=7/6 GFP males/females). **B)** Usage comparison for each behavioral syllable identified by MoSeq between DMS-M4- KO and GFP mice, organized by broad behavioral category. All syllables that collectively account for 99% of syllable usage across all recordings. Significance for each syllable determined by independent Mann-Whitney test. **C)** Total usage of all Grooming related syllables comparing M4 deletion and sex (2-way ANOVA p=0.1389 and p=0.0132) with a trend towards an interaction effect (p=0.0786). DMS M4 deletion increased grooming in males (Fisher’s LSD, p=0.0109) and female animals injected with GFP exhibited increased grooming compared to males injected with GFP (Fisher’s LSD, p=0.0017). **D)** Total usage of all active behaviors comparing M4 deletion and sex (2-way ANOVA, p=0.0621 and p=0.0077). DMS M4 deletion decreased active behaviors in males (Fisher’s LSD, p=0.0112) and female animals injected with GFP exhibited decreased grooming compared to males injected with GFP (Fisher’s LSD, p=0.0024). **E)** Total usage of all passive behaviors comparing M4 deletion and sex (2-way ANOVA p=0.7398 and p=0.6442).

### Selective deletion of M4 receptors from the DMS of SAPAP3-/- male mice does not inhibit VU’154 efficacy

SAPAP3-/- DMS-M4-KO mice were tested in the same VU’154 dose- response experiments as the SAPAP3-/- mice (**Fig. 5A**). SAPAP3-/- DMS-M4-KO male animals that went through full dose-response (n=7) showed an effect of VU’154 (Friedman test, p=0.0079; **Fig. 5B**). Active behaviors were not affected by VU’154 (Friedman test, p=0.1202; **Fig. 5C**), but Passive behaviors were (Friedman test, p=0.0182; **Fig. 5D**). Collectively, these results suggest that while deletion of M4 from the DMS can increase grooming in SAPAP3-/- mice, these receptors are not necessary for M4 PAMs to suppress repetitive grooming behaviors.

**Figure 5.**
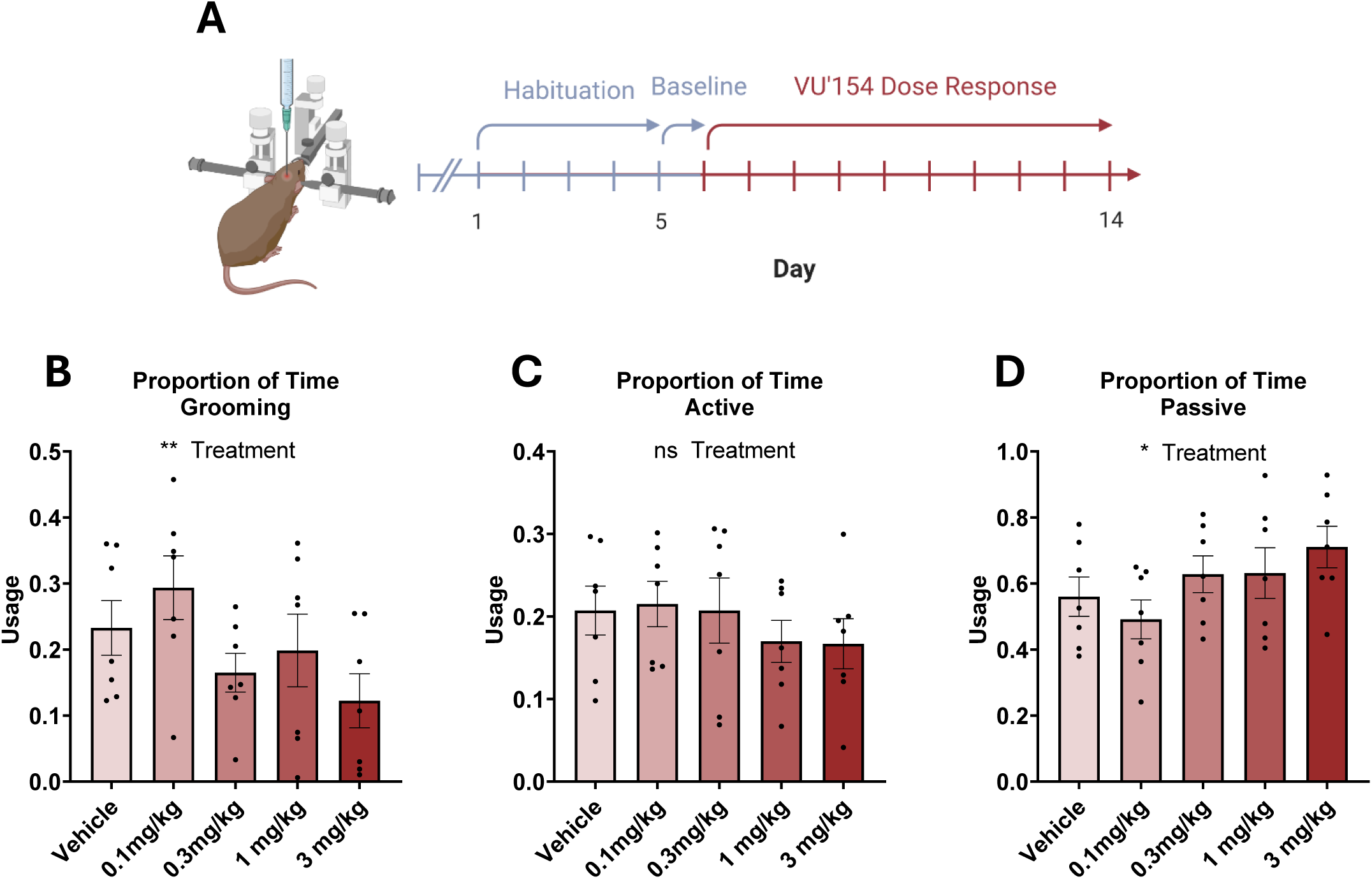
Behavioral analysis of SAPAP3^-/-^ DMS-M4-KO males in response to VU’154. **A)** 30 min recordings taken after 4 days of habituation and baseline recording. Recordings were taken every other day with a counterbalanced dosage schedule used where each animal received each dose once. **B)** VU’154 affects grooming in SAPAP3^-/-^ DMS-M4-KO males (Friedman test p=0.0079). **C)** VU’154 does not affect active behaviors in SAPAP3^-/-^ DMS M4KO males (Friedman test p=0.1202). **D)** VU’154 does affect passive behaviors in SAPAP3^-/-^ DMS M4KO (Friedman test p=0.0182).

## Discussion

Our results show that the M4 PAM VU’154 reduces excessive grooming in male SAPAP3-/- mice at an intermediate dose (0.3 mg/kg) that does not broadly reduce naturalistic, non- grooming behaviors. At the highest dose tested (3 mg/kg), behavioral output, including locomotion, was broadly reduced in both sexes and genotypes, indicating a therapeutic window in which VU’154 can reduce repetitive grooming relatively selectively in male mice. Surprisingly, VU’154 had little to no effect on repetitive grooming in female SAPAP3-/- mice, despite SAPAP3-/- mice showing no significant sex differences in baseline grooming. Additionally, deleting M4 receptors in the dorsomedial striatum (DMS) increased baseline grooming in male SAPAP3-/- x M4fl/fl but, surprisingly, did not eliminate the grooming- reducing effect of VU’154. Together, these findings suggest that M4 receptors contribute to the regulation of the excessive grooming observed in SAPAP3-/- mice, with sex-dependent effects and a distinction between the receptor populations regulating baseline grooming and those required for the behavioral response to systemic VU’154 administration.

The MoSeq characterization of SAPAP3-/- mice identified behavioral changes consistent with previous reports, including excessive grooming, increased inactivity, and decreased locomotion (Hadjas et al., 2020; Lamothe et al., 2023; Welch et al., 2007). We also report decreased rearing in SAPAP3-/- mice compared with littermate controls. Although we detected no sex differences in SAPAP3-/- mice under baseline conditions, we did observe significant sex differences in the grooming behavior of SAPAP3-/- x M4fl/fl mice, with female mice of this strain demonstrating a greater grooming phenotype than males, suggesting that slight changes in background strain may play a role in whether sex-specific changes are observed. Previous work has reported sexually dimorphic grooming in young-adult (5- 11-week-old) SAPAP3-/- mice (Wilson et al., 2025), while studies in 8-month-old mice show no significant sex differences in either grooming or somatosensory gating (Manning, Wang, Saikali, Winner, & Ahmari, 2021). Collectively, these findings suggest that sex-based differences in SAPAP3-/- behaviors may vary by developmental time point and background strain.

The sex-based differences in basal grooming make it challenging to interpret the finding that deleting M4 from the DMS increased grooming in male but not female mice. The grooming phenotype in female SAPAP3-/- x M4fl/fl mice may have been limited by a ceiling effect that obscured further increases in grooming. Alternatively, female SAPAP3-/- grooming may be modulated through partially different brain circuits than in males.

Ultimately, further studies assessing deletion of M4 from the dorsolateral striatum or nucleus accumbens are needed to determine if the baseline grooming phenotype is mediated via similar or distinct circuits in male and female SAPAP3-/- mice. The ability of VU’154 to reduce excessive grooming in male, but not female, SAPAP3-/- mice was unexpected. Previous work has demonstrated that the antipsychotic-like efficacy of VU’154 is observed in both male and female mice (Bubser et al., 2014; Joyner, Johnston, Choi, May, Carden, & Foster, 2026), and clinical studies have found no clear sex differences in efficacy of muscarinic-targeted therapies in patients with schizophrenia (Kaul et al., 2024). This divergence could suggest that the circuits or receptor populations through which VU’154 mediates antipsychotic-like effects may differ from those required for its effects on repetitive grooming. Previous studies have shown that the antipsychotic-like effects are mediated primarily via M4 receptors expressed on D1-expressing spiny projection neurons (Jeon et al., 2010; Joyner et al., 2026), and future studies are needed to determine whether these same receptors are necessary for VU’154-mediated reductions in repetitive grooming. Alternatively, SAPAP3 deficiency is associated with altered striatal ACh release (Malgady et al., 2023), as well as disrupted corticostriatal signaling (Wan et al., 2014), raising the possibility that M4-mediated reductions in grooming could be mediated in part by M4 heteroreceptors on glutamatergic terminals (Pancani et al., 2014), or M4 autoreceptors located on cholinergic interneurons (Gonzales et al., 2013). Interestingly, psilocybin has also been reported to reduce repetitive grooming in male but not female SAPAP3-/- mice (Gattuso, Wilson, Hannan, & Renoir, 2025). Collectively, these studies indicating sex-dependent drug efficacy point to the possibility of sex-dependent circuit changes underlying repetitive grooming in male vs. female mice. Further studies will be needed to rigorously test potential sex-based circuit changes to help elucidate the therapeutic potential of novel strategies for reducing repetitive behaviors using cholinergic- and serotonergic-based strategies.

Several sex-based differences in striatal biology have previously been reported, including a higher density of cholinergic interneurons in the striatum of male mice (Van Zandt, Flanagan, & Pittenger, 2024), disruptions in circadian locomotor activity in female but not male M4 KO mice (Valuskova, Forczek, Farar, & Myslivecek, 2018), and sex-specific differences in striatal dopamine release (Zachry, Nolan, Brady, Kelly, Siciliano, & Calipari, 2021). Our M4 RNA expression results showed that female animals exhibited more Chrm4 RNA puncta per cell than males, with no change in the total number of M4-expressing cells. This result was not unique to M4, as the number of Chrm1 RNA puncta per cell was also significantly greater in the striatum of females compared to males. These results are consistent with findings that female rats have elevated mAChR expression compared with male rats, as assessed using 3H-QNB binding assays (Miller, 1983). If M4 receptor levels are increased in the striatum of female mice, this could increase receptor reserve, producing a leftward shift in the ability of ACh to activate M4 receptors. Because PAMs act by potentiating endogenous ACh, females with greater receptor reserve could be closer to maximal M4 signaling at baseline, leaving less room for potentiation. Further studies will be needed to assess this possibility, including studies using autoradiography approaches combined with mAChR KO mice (Valuskova, Farar, Forczek, Krizova, & Myslivecek, 2018) in male and female mice, to rigorously assess whether sex-based changes in M1 and M4 protein levels match our sex-specific RNA results.

Though sex-specific differences in M4 expression levels may explain the ability of VU’154 to cause a sex-specific reduction in grooming in male mice, several other possibilities exist, including sex hormones. One limitation of the current study is that we did not measure estrous stage or circulating hormones, preventing direct assessment of their contribution. Although the distribution of grooming responses to VU’154 in female SAPAP3-/- mice did not exhibit an apparent bimodal or multimodal pattern, it is possible that M4 PAM efficacy was estrous cycle-dependent, and future studies will be needed to assess this possibility.

Collectively, these studies suggest that M4 PAMs may represent a novel therapeutic strategy for reducing repetitive behaviors that are observed across numerous disorders. Our results demonstrate that VU’154 can reduce repetitive grooming in male SAPAP3-/- mice in a dose-dependent manner, with efficacy at intermediate doses showing relatively specific reductions in repetitive grooming. Further studies are needed to identify the circuits through which VU’154 mediates reductions in repetitive grooming and to elucidate the mechanisms underlying the sex-dependent effects.

## Author Contributions

**CMJ:** Conceptualization, Methodology, Investigation, Data Analysis, Manuscript Writing and Editing. **DGJ**: Methodology, Investigation, Data Analysis, Manuscript Writing and Editing. **YC:** Methodology, Investigation, Manuscript Editing. **RDC:** Investigation, Manuscript Editing. **MNM:** Investigation, Manuscript Editing. **MW:** Investigation, Manuscript Editing. **PM:** Investigation and Manuscript Editing. **LB:** Investigation and Manuscript Editing. **KM:** Investigation and Manuscript Editing. **VP:** Investigation and Manuscript Editing. **CC:** Investigation and Manuscript Editing. **CO:** Methodology, Manuscript Editing. **DJF:** Conceptualization, Methodology, Resources, Supervision, Manuscript Writing and Editing.

## AI declaration

During manuscript preparation, the authors used ChatGPT (OpenAI) to assist with editing (Sol 6.1) and with code (Sol 5.X). The authors reviewed and revised all AI- assisted text and take responsibility for the final manuscript.

## Funding

This research was funded by the National Institute of Mental Health (R01MH122545 to DJF)

## Competing Interests

All authors declare no competing interests.

## Notes

### Competing Interest Statement

The authors have declared no competing interest.

## References

Ahmari, S. E., Spellman, T., Douglass, N. L., Kheirbek, M. A., Simpson, H. B., Deisseroth, K., … Hen, R. (2013). Repeated cortico-striatal stimulation generates persistent OCD- like behavior. Science, 340(6137), 1234–1239. doi:10.1126/science.1234733

Berezovskaia, A., Lindsley, C., Fink-Jensen, A., & Wortwein, G. (2025). M(4) Positive Allosteric Modulator VU0467154 Impacts Amphetamine Sensitization and Spontaneous Locomotion in Male Mice. ACS Chem Neurosci, 16(5), 868–879. doi:10.1021/acschemneuro.4c00795

Bubser, M., Bridges, T. M., Dencker, D., Gould, R. W., Grannan, M., Noetzel, M. J., … Jones, C. K. (2014). Selective activation of M4 muscarinic acetylcholine receptors reverses MK-801-induced behavioral impairments and enhances associative learning in rodents. ACS Chem Neurosci, 5(10), 920–942. doi:10.1021/cn500128b

Burguiere, E., Monteiro, P., Feng, G., & Graybiel, A. M. (2013). Optogenetic stimulation of lateral orbitofronto-striatal pathway suppresses compulsive behaviors. Science, 340(6137), 1243–1246. doi:10.1126/science.1232380

Burton, C. L., Longaretti, A., Zlatanovic, A., Gomes, G. M., & Tonini, R. (2024). Striatal insights: a cellular and molecular perspective on repetitive behaviors in pathology. Front Cell Neurosci, 18, 1386715. doi:10.3389/fncel.2024.1386715

Cadeddu, R., Braccagni, G., Branca, C., van Luik, E. R., Pittenger, C., Thomsen, M. S., & Bortolato, M. (2024). Activation of M(4) muscarinic receptors in the striatum reduces tic-like behaviours in two distinct murine models of Tourette syndrome. Br J Pharmacol, 181(17), 3064–3081. doi:10.1111/bph.16392

Gattuso, J. J., Wilson, C., Hannan, A. J., & Renoir, T. (2025). Psilocybin reduces grooming in the SAPAP3 knockout mouse model of compulsive behaviour. Neuropharmacology, 262, 110202. doi:10.1016/j.neuropharm.2024.110202

Gonzales, K. K., Pare, J. F., Wichmann, T., & Smith, Y. (2013). GABAergic inputs from direct and indirect striatal projection neurons onto cholinergic interneurons in the primate putamen. J Comp Neurol, 521(11), 2502–2522. doi:10.1002/cne.23295

Hadjas, L. C., Schartner, M. M., Cand, J., Creed, M. C., Pascoli, V., Luscher, C., & Simmler, L. D. (2020). Projection-specific deficits in synaptic transmission in adult Sapap3- knockout mice. Neuropsychopharmacology, 45(12), 2020–2029. doi:10.1038/s41386-020-0747-3

Jeon, J., Dencker, D., Wortwein, G., Woldbye, D. P., Cui, Y., Davis, A. A., … Wess, J. (2010). A subpopulation of neuronal M4 muscarinic acetylcholine receptors plays a critical role in modulating dopamine-dependent behaviors. J Neurosci, 30(6), 2396–2405. doi:10.1523/JNEUROSCI.3843-09.2010

Joyner, D. G., Johnston, C. M., Choi, Y., May, M. N., Carden, R. D., & Foster, D. J. (2026). Striatal M4 Receptors Mediate M4 PAM Antipsychotic-like Efficacy Independently of Normalizing Striatal Dopamine Release. bioRxivdoi:10.64898/2026.10.26.761976

Kaul, I., Sawchak, S., Claxton, A., Sauder, C., Hassman, H. H., Kakar, R., … Brannan, S. K. (2024). Efficacy of xanomeline and trospium chloride in schizophrenia: pooled results from three 5-week, randomized, double-blind, placebo-controlled, EMERGENT trials. Schizophrenia (Heidelb), 10(1), 102. doi:10.1038/s41537-024-00525-6

Kim, H., Lim, C. S., & Kaang, B. K. (2016). Neuronal mechanisms and circuits underlying repetitive behaviors in mouse models of autism spectrum disorder. Behav Brain Funct, 12(1), 3. doi:10.1186/s12993-016-0087-y

Lamothe, H., Schreiweis, C., Mondragon-Gonzalez, L. S., Rebbah, S., Lavielle, O., Mallet, L., & Burguiere, E. (2023). The Sapap3(-/-) mouse reconsidered as a comorbid model expressing a spectrum of pathological repetitive behaviours. Transl Psychiatry, 13(1), 26. doi:10.1038/s41398-023-02323-7

Langen, M., Bos, D., Noordermeer, S. D., Nederveen, H., van Engeland, H., & Durston, S. (2014). Changes in the development of striatum are involved in repetitive behavior in autism. Biol Psychiatry, 76(5), 405–411. doi:10.1016/j.biopsych.2013.08.013

Lin, S., Gillis, W. F., Weinreb, C., Zeine, A., Jones, S. C., Robinson, E. M., … Datta, S. R. (2024). Characterizing the structure of mouse behavior using Motion Sequencing. Nat Protoc, 19(11), 3242–3291. doi:10.1038/s41596-024-01015-w

Longo, F., Aryal, S., Anastasiades, P. G., Maltese, M., Baimel, C., Albanese, F., … Klann, E. (2023). Cell-type-specific disruption of cortico-striatal circuitry drives repetitive patterns of behavior in fragile X syndrome model mice. Cell Rep, 42(8), 112901. doi:10.1016/j.celrep.2023.112901

Malgady, J. M., Baez, A., Hobel, Z. B., Jimenez, K., Goldfried, J., Prager, E. M., … Plotkin, J. L. (2023). Pathway-specific alterations in striatal excitability and cholinergic modulation in a SAPAP3 mouse model of compulsive motor behavior. Cell Rep, 42(11), 113384. doi:10.1016/j.celrep.2023.113384

Maneeton, P., Maneeton, B., Winichaikul, Y., Kawilapat, S., Kienngam, N., & Maneeton, N. (2024). Updated Systematic Review and Meta-Analysis of Randomized Controlled Trials: Antidepressants for Restricted and Repetitive Behaviors in Autism Spectrum Disorder. Neuropsychiatr Dis Treat, 20, 1711–1723. doi:10.2147/NDT.S465611

Manning, E. E., Wang, A. Y., Saikali, L. M., Winner, A. S., & Ahmari, S. E. (2021). Disruption of prepulse inhibition is associated with compulsive behavior severity and nucleus accumbens dopamine receptor changes in Sapap3 knockout mice. Sci Rep, 11(1), 9442. doi:10.1038/s41598-021-88769-5

Miller, J. C. (1983). Sex differences in dopaminergic and cholinergic activity and function in the nigro-striatal system of the rat. Psychoneuroendocrinology, 8(2), 225–236. doi:10.1016/0306-4530(83)90059-8

O’Loghlen, J., McKenzie, M., Lang, C., & Paynter, J. (2025). Repetitive Behaviors in Autism and Obsessive-Compulsive Disorder: A Systematic Review. J Autism Dev Disord, 55(7), 2307–2321. doi:10.1007/s10803-024-06357-8

Pancani, T., Bolarinwa, C., Smith, Y., Lindsley, C. W., Conn, P. J., & Xiang, Z. (2014). M4 mAChR-mediated modulation of glutamatergic transmission at corticostriatal synapses. ACS Chem Neurosci, 5(4), 318–324. doi:10.1021/cn500003z

Piantadosi, S. C., Manning, E. E., Chamberlain, B. L., Hyde, J., LaPalombara, Z., Bannon, N. M., … Ahmari, S. E. (2024). Hyperactivity of indirect pathway-projecting spiny projection neurons promotes compulsive behavior. Nat Commun, 15(1), 4434. doi:10.1038/s41467-024-48331-z

Ramirez-Armenta, K. I., Alatriste-Leon, H., Verma-Rodriguez, A. K., Llanos-Moreno, A., Ramirez-Jarquin, J. O., & Tecuapetla, F. (2022). Optogenetic inhibition of indirect pathway neurons in the dorsomedial striatum reduces excessive grooming in Sapap3-knockout mice. Neuropsychopharmacology, 47(2), 477–487. doi:10.1038/s41386-021-01161-9

Robbins, T. W., Banca, P., & Belin, D. (2024). From compulsivity to compulsion: the neural basis of compulsive disorders. Nat Rev Neurosci, 25(5), 313–333. doi:10.1038/s41583-024-00807-z

Ruscio, A. M., Stein, D. J., Chiu, W. T., & Kessler, R. C. (2010). The epidemiology of obsessive-compulsive disorder in the National Comorbidity Survey Replication. Mol Psychiatry, 15(1), 53–63. doi:10.1038/mp.2008.94

Saxena, S., & Rauch, S. L. (2000). Functional neuroimaging and the neuroanatomy of obsessive-compulsive disorder. Psychiatr Clin North Am, 23(3), 563–586. doi:10.1016/s0193-953x(05)70181-7

Skapinakis, P., Caldwell, D. M., Hollingworth, W., Bryden, P., Fineberg, N. A., Salkovskis, P., … Lewis, G. (2016). Pharmacological and psychotherapeutic interventions for management of obsessive-compulsive disorder in adults: a systematic review and network meta-analysis. Lancet Psychiatry, 3(8), 730–739. doi:10.1016/S2215-0366(16)30069-4

Song, B. J., Ge, Y., Nicolella, A., Kwon, M. J., Lodder, B., Bonanno, K., … Sheng, M. (2025). Elevated synaptic PKA activity and abnormal striatal dopamine signaling in Akap11 mutant mice, a genetic model of schizophrenia and bipolar disorder. bioRxivdoi:10.1101/2024.09.24.614783

Steenfeldt-Kristensen, C., Jones, C. A., & Richards, C. (2020). The Prevalence of Self- injurious Behaviour in Autism: A Meta-analytic Study. J Autism Dev Disord, 50(11), 3857–3873. doi:10.1007/s10803-020-04443-1

Stein, D. J., Costa, D. L. C., Lochner, C., Miguel, E. C., Reddy, Y. C. J., Shavitt, R. G., … Simpson, H. B. (2019). Obsessive-compulsive disorder. Nat Rev Dis Primers, 5(1), 52. doi:10.1038/s41572-019-0102-3

Ting, J. T., & Feng, G. (2011). Neurobiology of obsessive-compulsive disorder: insights into neural circuitry dysfunction through mouse genetics. Curr Opin Neurobiol, 21(6), 842–848. doi:10.1016/j.conb.2011.04.010

Valuskova, P., Forczek, S. T., Farar, V., & Myslivecek, J. (2018). The deletion of M(4) muscarinic receptors increases motor activity in females in the dark phase. Brain Behav, 8(8), e01057. doi:10.1002/brb3.1057

Van Zandt, M., Flanagan, D., & Pittenger, C. (2024). Sex differences in the distribution and density of regulatory interneurons in the striatum. Front Cell Neurosci, 18, 1415015. doi:10.3389/fncel.2024.1415015

Vuckovic, Z., Wang, J., Pham, V., Mobbs, J. I., Belousoff, M. J., Bhattarai, A., … Thal, D. M. (2023). Pharmacological hallmarks of allostery at the M4 muscarinic receptor elucidated through structure and dynamics. Elife, *12*doi:10.7554/eLife.83477

Wan, Y., Ade, K. K., Caffall, Z., Ilcim Ozlu, M., Eroglu, C., Feng, G., & Calakos, N. (2014). Circuit-selective striatal synaptic dysfunction in the Sapap3 knockout mouse model of obsessive-compulsive disorder. Biol Psychiatry, 75(8), 623–630. doi:10.1016/j.biopsych.2013.01.008

Welch, J. M., Lu, J., Rodriguiz, R. M., Trotta, N. C., Peca, J., Ding, J. D., … Feng, G. (2007). Cortico-striatal synaptic defects and OCD-like behaviours in Sapap3-mutant mice. Nature, 448(7156), 894–900. doi:10.1038/nature06104

Welch, J. M., Wang, D., & Feng, G. (2004). Differential mRNA expression and protein localization of the SAP90/PSD-95-associated proteins (SAPAPs) in the nervous system of the mouse. J Comp Neurol, 472(1), 24–39. doi:10.1002/cne.20060

Wilson, C., Gattuso, J. J., Kuznetsova, M., Li, S., Ang, C. S., Williamson, N. A., … Renoir, T. (2025). Early-life social isolation stress worsens compulsive- and anxiety-like behaviour and alters gene expression in the prefrontal cortex of SAPAP3 knockout mice. J Affect Disord, 388, 119600. doi:10.1016/j.jad.2025.119600

Wiltschko, A. B., Johnson, M. J., Iurilli, G., Peterson, R. E., Katon, J. M., Pashkovski, S. L., … Datta, S. R. (2015). Mapping Sub-Second Structure in Mouse Behavior. Neuron, 88(6), 1121–1135. doi:10.1016/j.neuron.2015.11.031

Wiltschko, A. B., Tsukahara, T., Zeine, A., Anyoha, R., Gillis, W. F., Markowitz, J. E., … Datta, S. R. (2020). Revealing the structure of pharmacobehavioral space through motion sequencing. Nat Neurosci, 23(11), 1433–1443. doi:10.1038/s41593-020-00706-3

Yu, Y., Chaulagain, A., Pedersen, S. A., Lydersen, S., Leventhal, B. L., Szatmari, P., … Skokauskas, N. (2020). Pharmacotherapy of restricted/repetitive behavior in autism spectrum disorder:a systematic review and meta-analysis. BMC Psychiatry, 20(1), 121. doi:10.1186/s12888-020-2477-9

Zachry, J. E., Nolan, S. O., Brady, L. J., Kelly, S. J., Siciliano, C. A., & Calipari, E. S. (2021). Sex differences in dopamine release regulation in the striatum. Neuropsychopharmacology, 46(3), 491–499. doi:10.1038/s41386-020-00915-1

Zeidan, J., Fombonne, E., Scorah, J., Ibrahim, A., Durkin, M. S., Saxena, S., … Elsabbagh, M. (2022). Global prevalence of autism: A systematic review update. Autism Res, 15(5), 778–790. doi:10.1002/aur.2696

Zhou, M. S., Nasir, M., Farhat, L. C., Kook, M., Artukoglu, B. B., & Bloch, M. H. (2021). Meta- analysis: Pharmacologic Treatment of Restricted and Repetitive Behaviors in Autism Spectrum Disorders. J Am Acad Child Adolesc Psychiatry, 60(1), 35–45. doi:10.1016/j.jaac.2020.03.007

