## Supplemental Materials for "M4 Muscarinic Receptor Positive Allosteric Modulation Reduces Repetitive Grooming in Male SAPAP3-/- Mice in a Sex-Dependent Manner"

### Supplemental Methods

**Animals** SAPAP3$^{-/-}$ mice were generously provided by Dr. Jeremy Veenstra-Vanderweele and were bred by crossing SAPAP3$^{+/-}$ mice to produce SAPAP3$^{-/-}$ and littermate SAPAP3$^{+/+}$ controls. To generate region-specific M4 deletions, M4$^{fl/fl}$ mice were crossed with SAPAP3$^{-/-}$ mice to M4$^{fl/fl}$ x SAPAP3$^{+/-}$ breeders, which produced both M4$^{fl/fl}$ x SAPAP3$^{-/-}$ and M4$^{fl/fl}$ x SAPAP3$^{+/+}$ controls (Jeon et al., 2010). The behavioral laboratory was climate controlled with temperatures being kept at 73 ± 3°F with humidity maintained at 50 ± 20%. For the remaining animals outliers were detected via ROUT with a Q of 1% in each behavioral category. Outliers in any category were removed from analysis in all categories, resulting in one exclusion. After applying these exclusion criteria, the final number was 35 animals. SAPAP3$^{-/-}$ n=18 (females n=9, males n=9), and WT control n = 17 (females n = 8, males n=9). For M4 deletion cohorts, animals were excluded if the M4 area of deletion was unilateral (excluded 3 animals) or below 15% of the DMS (excluded 0 animals) or exceeded 15% of the NAc or DLS (excluded 3 animals).

**Recording Setup** Two or three open field tanks and cameras were used in parallel with animals recorded at the same time being selected from the same home cage where possible.

**Behavioral Recordings** Animals were brought to a behavioral laboratory in their home cages at the start of the light cycle. Animals were left in their home cages to habituate to the room for 30 minutes prior to testing. On days in which animals were habituated to open field they were given an i.p. injection of 10 mL/kg sterile saline and placed into the open field tanks for 10 minutes without recording. Mice were then removed, and the tanks were cleaned with 70% ethanol and allowed to dry prior to more mice being habituated. Mice were not injected prior to 30-minute baseline recordings. During pharmacological dose-response experimental recordings mice were injected with 10 mL/kg of either vehicle or VU'154 and returned to their home cage for 15 minutes prior to 30-minute recordings in the open field. Temperature and Humidity were logged at the beginning of each recording session.

**MoSeq Data Processing** The SAPAP3$^{-/-}$ dose response model was trained using vehicle and 1 mg/kg VU'154 recordings from the SAPAP3$^{-/-}$ animals. The SAPAP3$^{-/-}$ DMS-M4-KO dose-response model was trained using vehicle and 1 mg/kg VU'154 recordings from SAPAP3$^{-/-}$ DMS-M4-KO animals. These training conditions were selected based on preliminary results suggesting a behavioral effect at 1 mg/kg and were retained across both dose response models for consistency.

**Manual Scoring of Grooming** Scorers were trained to record grooming start and stop times in 30-minute videos. Training was performed using example videos to assess inter-scorer reliability. F1 values of 0.85 or greater between scorers on training videos were used as a cut-off for selecting scorers for experimental recordings. Scorers were blinded to experimental conditions. Baseline recordings of experiments classifying the SAPAP3$^{-/-}$ phenotype were scored fully. Other models used custom code to randomly select 2-minute clips from each video analyzed to quantify model efficacy.

**Locomotion and Rearing Detection** Scalar values collected via MoSeq's data pipeline was used to separately determine the usage of locomotion and rearing separate from individual syllables. Velocity values for detected mouse centroids were averaged across consecutive three-frame bins to generate a nominal 10-Hz time series. Locomotion was calculated as the proportion of all bins in the recording with a mean velocity ≥ 4cm/s (Jordan & Keller, 2020). Rearing was calculated based on a per mouse height threshold. The average-height values were divided into 10 equal-width bins, and the mean height within the most populated bin was used as an estimate of the predominant low-height posture. A normal distribution was fitted by maximum likelihood to height values at or below this estimate, accounting for upper truncation of that value. The rearing threshold was defined as the fitted mean plus five fitted standard deviations. Rearing usage was calculated as the proportion of all recorded frames with an average height at or above this threshold.

**RNAscope** Mice were anesthetized in 2-3% isoflurane and euthanized by decapitation. Brains were rapidly removed, flash-frozen in 2-methylbutane, and stored at -80°C. Brains were subsequently sectioned coronally (20 µm) on a cryostat, collected on SuperFrost Plus slides (Fisher) and stored in a slide box that was pretreated with RNase Zap (Fisher). Before the assay, slices were fixed for 1 h at 4°C in pre-chilled 4% PFA prepared in RNase-free PBS and then rinsed twice in RNase-free PBS. Sections were then dehydrated sequentially in 50%, 70%, 100%, and 100% ethanol. Probe mixtures were prepared according to the number of targets to be labeled using a 1x diluted probe mixture for Channel 1 as diluent as follows: M1 was labeled on Channel 1 (Mm-Chrm1-O1, 96%, ACD 483441), M4 on Channel 2 (Mm-Chrm4-C2, 2%, ACD 410581-C2), and ChAT on Channel 3 (Mm-Chat-C3, 2% ACD 408731-C3). Fluorophores used were Channel 1: TSA Vivid 570 (yellow, 1:4000), Channel 2: TSA Vivid 650 (red, 1:5000), and Channel 3: TSA Vivid 520 (green, 1:3000). The assay was performed according to the manufacturer’s instructions. Following the multiplex assay, sections were incubated in DAPI (RNAscope Multiplex V2 DAPI) for 30 s, excess was removed, and sections were mounted with ProLong Gold Antifade Mountant (Fisher) and cover slipped. Sections were allowed to dry for at least 30 minutes and stored in the dark at 4 °C. Slides were imaged on the Leica Stellaris 5 LIAchroic Confocal System within 1 week of the assay.

**RNAscope image analysis** To measure the ratio of M1- and M4-positive cells to total cells, M4, M1, and DAPI channels were converted to Luminance, then to RGB respectively in Fiji (Image J), and then analyzed in QuPath. QuPath’s Cell Detection was run on a non-deleted area (dorsolateral striatum) with default values except for the following changes: Background Radius: 10 µm; Median Filter Radius: 2 µm; Sigma: 2 µm; Threshold: 1. Custom code optimized M1 and M4 puncta detection thresholds using the non-deleted image (https://doi.org/10.5281/zenodo.23301431), and these thresholds were utilized for puncta detection in the deleted area (dorsomedial striatum). Afterward, the area of deletion was determined utilizing Fiji’s BigWarp plugin to fit fluorescent images to the corresponding atlas image and outlining the deletion.

### References

Jeon, J., Dencker, D., Wortwein, G., Woldbye, D. P., Cui, Y., Davis, A. A., … Wess, J. (2010). A subpopulation of neuronal M4 muscarinic acetylcholine receptors plays a critical role in modulating dopamine-dependent behaviors. *J Neurosci, 30*(6), 2396-2405. doi:10.1523/JNEUROSCI.3843-09.2010

Jordan, R., & Keller, G. B. (2020). Opposing Influence of Top-down and Bottom-up Input on Excitatory Layer 2/3 Neurons in Mouse Primary Visual Cortex. *Neuron, 108*(6), 1194-1206 e1195. doi:10.1016/j.neuron.2020.09.024

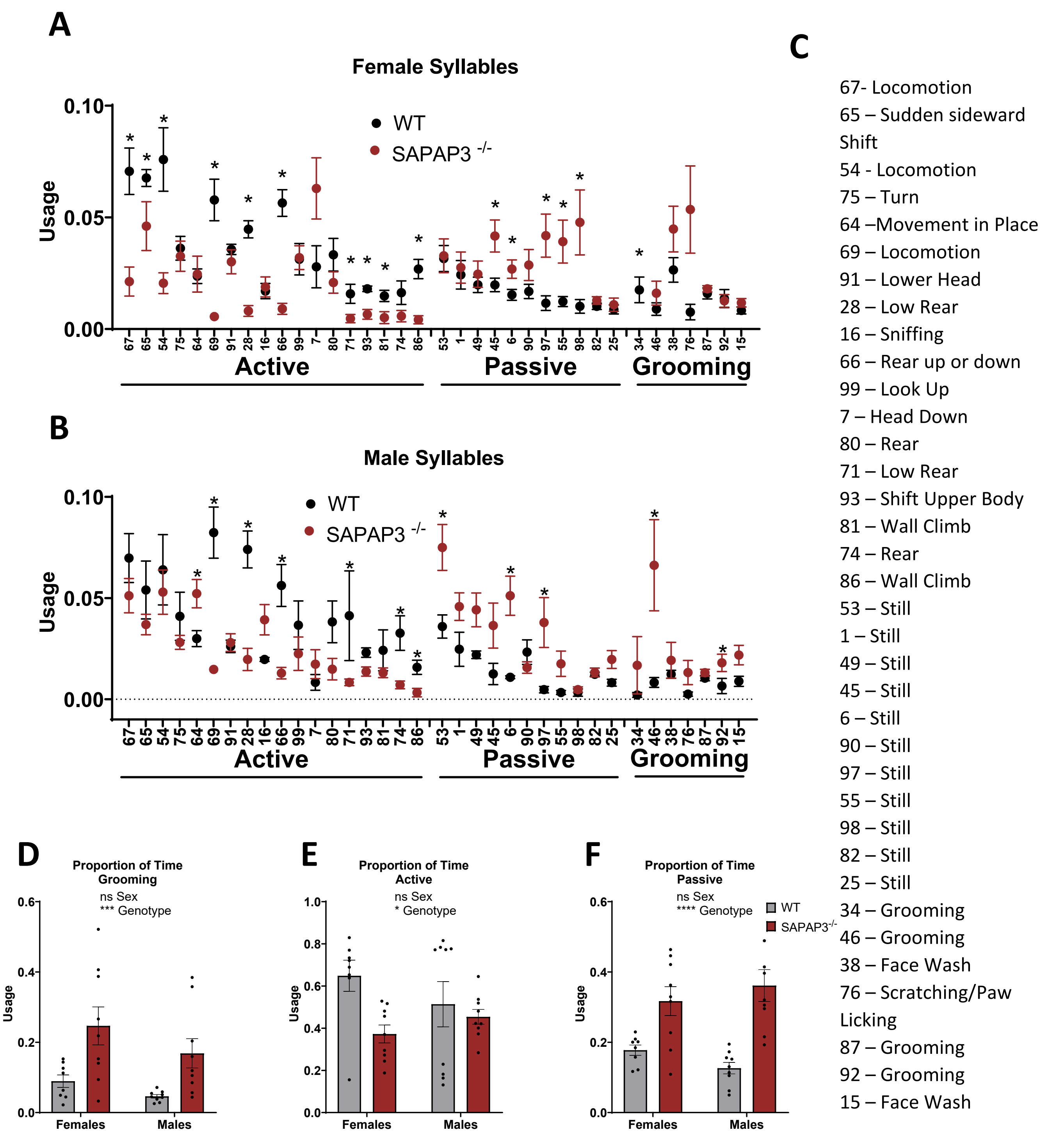

**Figure S1.** MoSeq analysis of SAPAP3$^{-/-}$ mice as compared to WT control littermates. Behavioral analysis of SAPAP3$^{-/-}$ mice as compared to WT control littermates showing no sex differences. **A-B)** Comparison of usage of each behavioral syllable identified by MoSeq, organized by broad behavioral category in SAPAP3$^{-/-}$ Female **(A)** and Male **(B)** mice. All syllables that collectively account for 95% of syllable usage across all recordings. **C)** Specific behaviors observed associated with each MoSeq identified syllable. **D)** Total usage of all Grooming related syllables comparing genotype and sex (2-way ANOVA p=0.0005 and p=0.1046). **E)** Total usage of all Active, non-grooming syllables comparing genotype and sex (2-way ANOVA p=0.0236 and p=0.7075). **F)** Total usage of all Passive syllables comparing genotype and sex (2-way ANOVA p<0.0001 and p=0.9174).

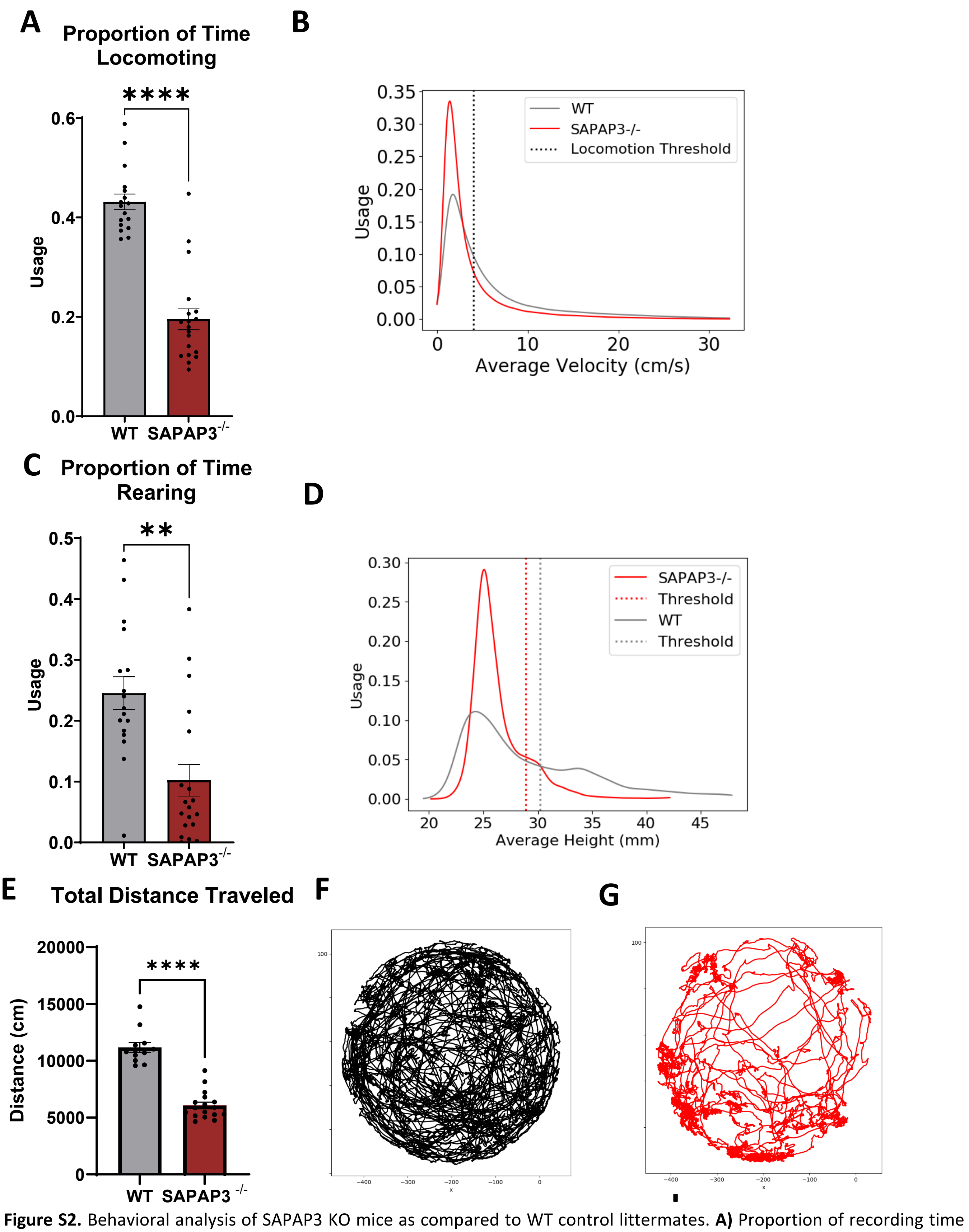

**Figure S2.** Behavioral analysis of SAPAP3 KO mice as compared to WT control littermates. **A)** Proportion of recording time spent locomoting between genotypes (Mann-Whitney test p<0.0001). **B)** Kernel density estimates of pooled, down-sampled velocities (3 frame means) for WT and SAPAP3$^{-/-}$ animals. Threshold for locomotion set at 4 cm/s. **C)** Proportion of time spent rearing between genotypes (Mann-Whitney test p=0.001). **D)** Kernel density estimates of pooled, down-sampled height (3 frame means) distributions from sample sessions selected for closely matched baseline height. Rearing detection threshold calculated as 5σ above normal distribution centered around mode height. **E)** Total distance traveled between genotypes (Mann-Whitney test p<0.0001). I) Proportion of time spent rearing between genotypes (Mann-Whitney test p=0.001). **F-G)** Example traces of animal movement over 30-minute recording selected from median distance traveled within **(F)** WT and **(G)** SAPAP3$^{-/-}$.

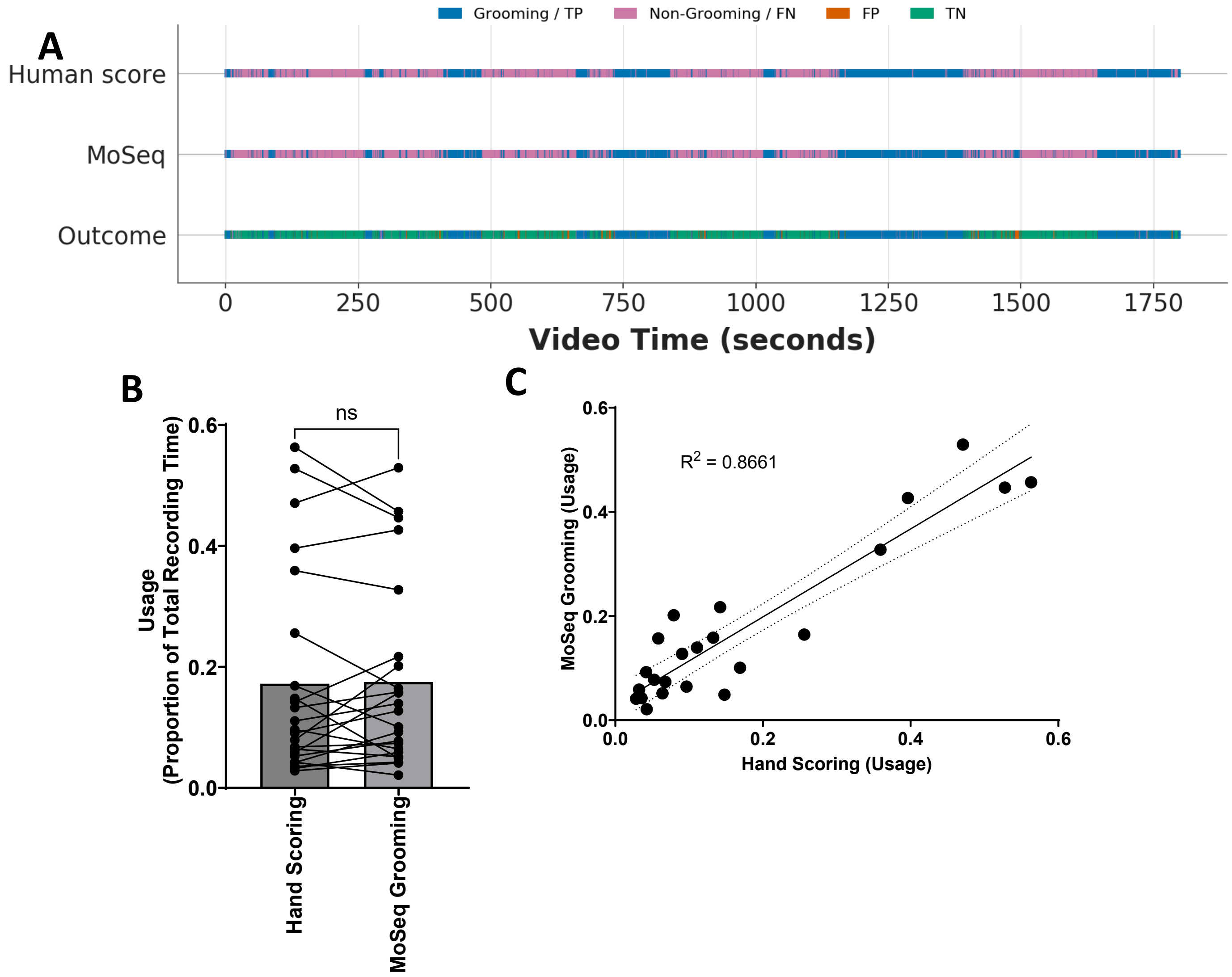

**Figure S3.** Validation of MoSeq Grooming Scoring. **A)** Example video grooming scores by blinded human scorer (top row) and MoSeq grooming associated syllables (middle row) with per frame Grooming (blue) and non-grooming (pink). Third row depicts MoSeq comparison to human scoring with per frame True Positives (blue), True Negatives (green), False Positives (orange), and False Negatives (pink). **B)** Total Grooming time during recordings as determined by human scorers and from summation of MoSeq Grooming associated syllables in 25 randomly selected videos. **C)** Correlation of total time spent grooming per recording between human and MoSeq scoring.

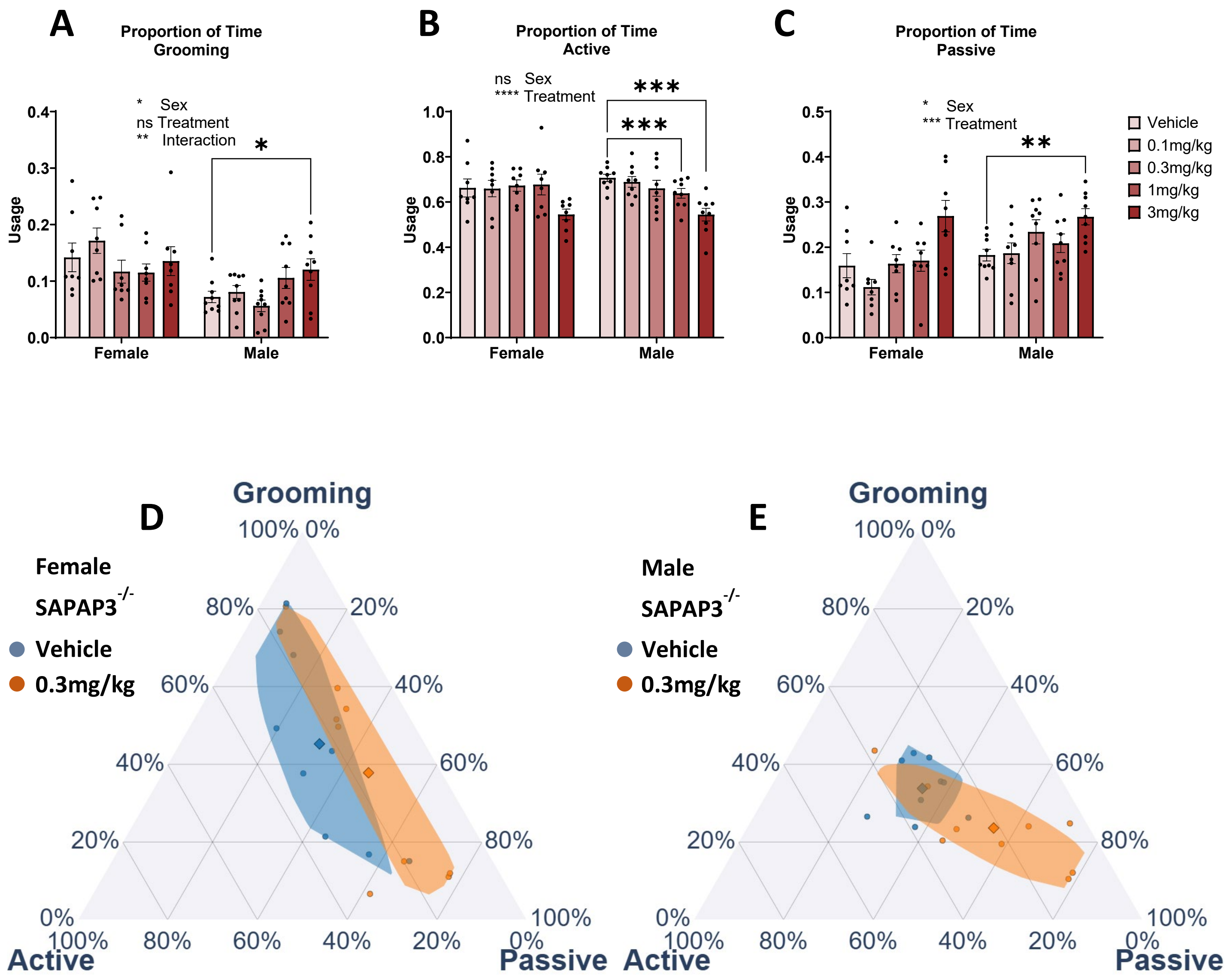

**Figure S4.** MoSeq analysis of WT dose response curve and SAPAP3$^{-/-}$ mice as compared to WT vehicle vs 3 mg/kg VU'154. **A)** Grooming behavior exhibited main effects of sex and sex X treatment interaction but not treatment in WT animals (2-way ANOVA, p=0.0156, p=0.0273, and p=0.0600). Increase in grooming observed in male but not female animals (Dunnett's post-hoc, p=0.0161 and p=0.9990). **B)** Active behaviors exhibited main treatment but not sex effects in WT animals (2-way ANOVA p<0.0001 and p=0.8969). Reductions exhibited at 1 mg/kg VU'154 and 3 mg/kg VU'154 in males (Dunnett's post-hoc, p=0.0009 and p=0.0003) but not in females (p=0.8577 and p=0.0944). **C)** Passive behaviors exhibited main effects in sex and treatment in WT animals (2-way ANOVA p=0.0358 and p=0.0003). Passive behavior increased at 3 mg/kg VU'154 in male but not female animals (Dunnett's post-hoc, p=0.0028, 0.2016). **D-E)** Ternary plot depicting per-recording behavioral usage across the 3 identified behavioral domains comparing Vehicle (blue) and 3 mg/kg dosage trials (orange) within subject in **D)** female SAPAP3$^{-/-}$ mice and **E)** male SAPAP3$^{-/-}$ mice

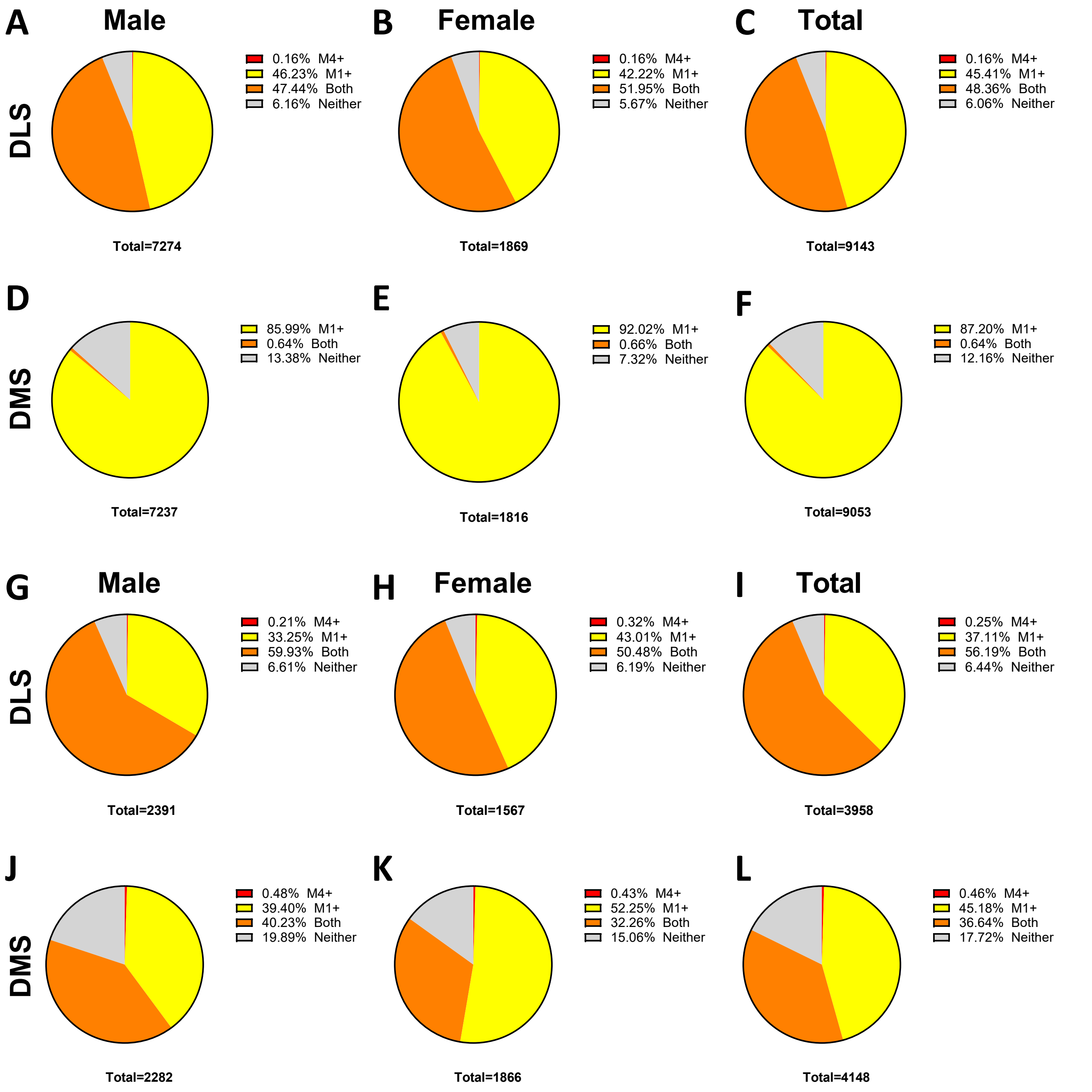

**Figure S5.** Comparative proportions of M4- and M1- positive cells in SAPAP3$^{-/-}$ DMS KO mice **(A-F)** split between males (n=7) **(A, D)** and females (n=3) **(B, E)**. Comparative proportions of M4- and M1- positive cells in SAPAP3$^{-/-}$ mice **(G-L)** split between males (n=4) **(D, J)** and females (n=3) **(H, K)**.

| | Figure(s) | Data | Statistics | Main Effect | | | Interaction | | |
|---|---|---|---|---|---|---|---|---|---|
| | | | | | F (DFn, DFd) | P value | | F (DFn, DFd) | Pvalue |
| SAPAP3-/- vs WT Dose Response | 2C, S4A | Grooming | 3-way ANOVA | Treatment | F (3.715, 115.2) = 3.070 | *P=0.0218 | Treatment x Genotype | F (4, 124) = 2.998 | *P=0.0211 |
| | | | | Genotype | F (1, 31) = 25.23 | ****P<0.0001 | Treatment x Sex | F (4, 124) = 1.691 | P=0.1563 |
| | | | | Sex | F (1, 31) = 6.143 | *P=0.0188 | Genotype x Sex | F (1, 31) = 0.7818 | P=0.3834 |
| | | | | | | | Treatment x Genotype x Sex | F (4, 124) = 0.9895 | P=0.4159 |
| | 2D, S4B | Active | 3-way ANOVA | Treatment | F (3.004, 93.11) = 21.75 | ****P<0.0001 | Treatment x Genotype | F (4, 124) = 0.7964 | P=0.5297 |
| | | | | Genotype | F (1, 31) = 269.5 | ****P<0.0001 | Treatment x Sex | F (4, 124) = 0.9443 | P=0.4408 |
| | | | | Sex | F (1, 31) = 1.924 | P=0.1753 | Genotype x Sex | F (1, 31) = 1.418 | P=0.2427 |
| | | | | | | | Treatment x Genotype x Sex | F (4, 124) = 1.573 | P=0.1855 |
| | 2E, S4C | Passive | 3-way ANOVA | Treatment | F (2.757, 85.46) = 13.00 | ****P<0.0001 | Treatment x Genotype | F (4, 124) = 0.6990 | P=0.5941 |
| | | | | Genotype | F (1, 31) = 23.90 | ****P<0.0001 | Treatment x Sex | F (4, 124) = 1.166 | P=0.3291 |
| | | | | Sex | F (1, 31) = 7.988 | **P=0.0082 | Genotype x Sex | F (1, 31) = 1.797 | P=0.1898 |
| | | | | | | | Treatment x Genotype x Sex | F (4, 124) = 0.8272 | P=0.5102 |

| | Figure(s) | Data | Statistics | Main Effect | | | Interaction | | |
|---|---|---|---|---|---|---|---|---|---|
| | | | | | F (DFn, DFd) | P value | | F (DFn, DFd) | Pvalue |
| Within SAPAP3-/- Dose Response | 2C | Grooming | 2-way ANOVA | Sex | F (1, 16) = 3.272 | P=0.0893 | Sex x Treatment | F (4, 64) = 0.8076 | P=0.5249 |
| | | | | Treatment | F (3.576, 57.22) = 3.303 | *P=0.0202 | | | |
| | 2D | Active | 2-way ANOVA | Sex | F (1, 16) = 3.689 | P=0.0728 | Sex x Treatment | F (4, 64) = 1.221 | P=0.3109 |
| | | | | Treatment | F (2.783, 44.53) = 8.185 | ***P=0.0003 | | | |
| | 2E | Passive | 2-way ANOVA | Sex | F (1, 16) = 5.120 | *P=0.0379 | Sex x Treatment | F (4, 64) = 0.9864 | P=0.4214 |
| | | | | Treatment | F (2.591, 41.46) = 6.391 | **P=0.0018 | | | |

| | Figure(s) | Data | Statistics | Main Effect | | | Interaction | | |
|---|---|---|---|---|---|---|---|---|---|
| | | | | | F (DFn, DFd) | P value | | F (DFn, DFd) | Pvalue |
| DMS-M4-KO Baseline Characterization | 4C | Grooming | 2-way ANOVA | Sex | F (1, 19) = 7.478 | *P=0.0132 | Sex x Deletion | F (1, 19) = 3.454 | P=0.0786 |
| | | | | Deletion | F (1, 19) = 2.387 | P=0.1389 | | | |
| | 4D | Active | 2-way ANOVA | Sex | F (1, 19) = 8.873 | **P=0.0077 | Sex x Deltion | F (1, 19) = 1.982 | P=0.1753 |
| | | | | Deletion | F (1, 19) = 3.929 | P=0.0621 | | | |
| | 4E | Passive | 2-way ANOVA | Sex | F (1, 19) = 0.2203 | P=0.6442 | Sex x Deletion | F (1, 19) = 0.4711 | P=0.5008 |
| | | | | Deletion | F (1, 19) = 0.1136 | P=0.7398 | | | |

| | Figure(s) | Data | Statistics | Main Effect | | | Interaction | | |
|---|---|---|---|---|---|---|---|---|---|
| | | | | | F (DFn, DFd) | P value | | F (DFn, DFd) | Pvalue |
| SAPAP3-/- vs WT Baseline Characterization | S1D | Grooming | 2-way ANOVA | Sex | F (1, 31) = 2.795 | P=0.1046 | Sex x Genotype | F (1, 31) = 0.2361 | P=0.6305 |
| | | | | Genotype | F (1, 31) = 14.86 | ***P=0.0005 | | | |
| | S1E | Active | 2-way ANOVA | Sex | F (1, 31) = 0.1434 | P=0.7075 | Sex x Genotype | F (1, 31) = 2.337 | P=0.1365 |
| | | | | Genotype | F (1, 31) = 5.665 | *P=0.0236 | | | |
| | S1F | Passive | 2-way ANOVA | Sex | F (1, 31) = 0.01093 | P=0.9174 | Sex x Genotype | F (1, 31) = 2.059 | P=0.1613 |
| | | | | Genotype | F (1, 31) = 31.62 | ****P<0.0001 | | | |

| | Figure(s) | Data | Statistics | Main Effect | | | Interaction | | |
|---|---|---|---|---|---|---|---|---|---|
| | | | | | F (DFn, DFd) | P value | | F (DFn, DFd) | Pvalue |
| Within WT Dose Response | S4A | Grooming | 2-way ANOVA | Sex | F (1, 15) = 7.432 | *P=0.0156 | Sex X Treatment | F (4, 60) = 2.947 | *P=0.0273 |
| | | | | Treatment | F (3.010, 45.15) = 2.650 | P=0.0600 | | | |
| | S4B | Active | 2-way ANOVA | Sex | F (1, 15) = 0.01736 | P=0.8969 | Sex X Treatment | F (4, 60) = 1.355 | P=0.2601 |
| | | | | Treatment | F (2.464, 36.96) = 16.11 | ****P<0.0001 | | | |
| | S4C | Passive | 2-way ANOVA | Sex | F (1, 15) = 5.316 | *P=0.0358 | Sex X Treatment | F (4, 60) = 1.135 | P=0.3487 |
| | | | | Treatment | F (2.574, 38.60) = 8.845 | ***P=0.0003 | | | |

**Table S1.** Main and interaction effects of all reported 2- and 3-way ANOVAs
